# Brain-derived circulating cell-free DNA defines the brain region and cell specific origins associated with neuronal atrophy

**DOI:** 10.1101/538827

**Authors:** Chatterton. Zac, Mendelev. Natalia, Chen. Sean, Raj. Towfique, Walker. Ruth, Carr. Walter, Kamimori. Gary, Beeri. Michal, Ge. Yongchao, Dwork. Andrew, Haghighi. Fatemeh

## Abstract

Liquid biopsies are revolutionizing the fields of prenatal non-invasive testing and cancer diagnosis by leveraging the genetic differences between mother and fetus, and, host and cancer. In the absence of genetic variance, epigenetics has been used to decipher the cell-of-origin of cell-free DNA (cfDNA). Liquid biopsies are minimally invasive and thus represent an attractive option for hard to biopsy tissues such as the brain. Here we report the first evidence of neuron derived cfDNA and cerebellum cfDNA within acute neurotrauma and chronic neurodegeneration, establishing the first class of peripheral biomarkers with specificity for the cell-type and brain-region undergoing potential injury and/or neurodegeneration.

## INTRODUCTION

It has been two decades since the first descriptions of fetal derived cell-free DNA (cfDNA) within maternal plasma [1] and the identification of tumor derived DNA within patient blood [2, 3]. Next generation sequencing (NGS) of these rare DNA fragments have established a minimally invasive technique (blood draw) for the diagnosis and disease tracking of cancer and graft rejection, and for non-invasive prenatal testing (NIPT). The technological advancements in NGS have increased the limits of detection of these techniques [4] to ~0.02%, which hold the promise of early detection of smaller cancers and more sensitive clinical patient tracking. However, in the absence of genetic markers DNA methylation has been used to delineate fetal from maternal DNA [5]. The use of cfDNA to identify fetal genetic abnormalities and to diagnose cancers, known as a liquid biopsies, largely removes the risk of serious fetal injury and the potential risks and difficulties of cancer biopsies [6]. This aspect is particularly attractive for neurological diagnosis. Liquid biopsies are capable of genotyping neurological tumors e.g. neuroblastoma by MYNC [7], and glioblastoma by EGFR mutations [8]. Fetal-derived cfDNA is detectable in mothers’ blood from 7 weeks of gestation when the fetus weighs < 1 gram [9]. Considering the average adult human brain weighs ~1.3kg and can undergo extensive tissue loss as a result of both neurotrauma and degeneration, it is likely that the apoptotic/necrotic DNA of the brain is shed into the peripheral blood. A seminal study identified the presence of brain-derived cfDNA in blood from subjects who had suffered a Traumatic Brain Injury (TBI), ischemic brain damage following cardiac arrest, and multiple sclerosis [10], providing the first evidence of brain-derived cfDNA during neurodegeneration.

DNA methylation is an epigenetic modification that is found covalently bound to cytosines within a CpG context. Uniquely to neurons, global DNA methylation is gained postnatally within the CpH context (H= any base other than guanine) [11], providing unique DNA methylation markers of neuronal cell identity. In light of the fact that DNA methylation is intricately involved in the regulation of gene expression it is likely that DNA methylation can not only delineate cell types but also the brain regions from which they are derived. Here we define CpG and CpH methylation that are specific to neurons of the dorsolateral prefrontal cortex (DLPFC-NeuN+), and CpG methylation with brain region specificity-in particular distinguishing the cerebellum from other brain regions. We developed a targeted multiplex next generation bisulfite sequencing (tNGBS) method assaying genomic regions of DLPFC-NeuN+ and cerebellum CpG/ CpH methylation, and perform ultra-deep sequencing (up to 100,000X) of cfDNA from human cohorts with possible indication of acute neurotrauma (blast wave exposure), during cognitive decline and with a chronic neurodegenerative disease (Parkinson’s Disease [PD]). Furthermore, we present new analytical methods for cell-of-origin deconvolution at single molecule resolution using k-mer analysis of raw bisulfite sequencing reads, establishing an ultra-fast and precise method for bisulfite sequencing based cell deconvolution (methylK). We identify an increase in DLPFC-NeuN+ cfDNA in response to blast wave exposure. Moreover, cognitive decline was associated with a decrease in DLPFC-NeuN+ cfDNA and evidence of cerebellum cfDNA. PD subjects exhibited reduced DLPFC-NeuN+ cfDNA across two independent cohorts.

Our platform provides a framework for an economical and scalable method for the identification of cell and brain-region specific neurodegeneration through the blood. Molecular biomarkers of injury and neurodegenerative disease are of critical importance in clinical research, both for the pre-symptomatic identification of vulnerable subjects, and as a marker of injury/ disease progression in clinical trials. These represent much needed new endpoints for clinical trials for diagnostic indicators of neurotrauma and prognostic indicators of progression of neurodegenerative disease.

## RESULTS

### Identification of brain-specific DNA methylation loci for targeted bisulfite sequencing

The majority of cfDNA within circulation is derived from blood cells. To identify CpG methylation loci capable of discriminating brain-derived DNA we implemented a 4-step pipeline (Figure 1A), initially performing linear modeling between CpG methylation microarray (Illumina HM450K) profiles from NeuN+ cells (n=71; DLPFC and OFC), NeuN-cells (n=78; DLPFC and OFC) and 63 profiles generated from brain tissue sections by the BrainSpan consortium (Supplementary Table 2) (Figure 1A). Each brain-tissue/cell was contrast with the CpG methylation microarray from 47 whole blood/PBMCs (Supplementary Table 2). We found a greater number of significant (adjusted P-value <0.05 & differential DNA methylation beta-value >50%) hypermethylated than hypomethylated CpG loci when comparing brain-tissue/cells and blood; notably neurons were found to have the greatest number of hypermethylated CpG (n=14048) (Figure 1B). We also contrast brain-tissue/cells with an additional 36 tissues/cell-types of the human body derived from the 3 germ layers (Supplementary Table 2) to refine genomic loci with brain-tissue/cell-specific CpG methylation. Again, we found a greater number of hypermethylated CpG loci within brain, particularly within neurons (Figure 1C). Using stricter thresholds for NeuN+ cells, we identified 45 CpG sites (Supplementary Table 11) that overlapped between the blood and lineage comparisons that exhibited >90% and 80% differential CpG methylation between neurons respectively (Figure 1D). Notably multiple methylated CpG loci were annotated to the Adenomatosis Polyposis Coli 2 (APC2) gene (3 CpG), Tripartite Motif Containing 38 (TRIM38) gene (2 CpG), SP100 Nuclear Antigen (SP100) gene (3 CpG) and the Transmembrane Protein 106A (TMEM106A) gene (5 CpG), in addition to 20 intergenic CpG loci. We identified 13 CpG sites that were >80% and 50% differentially methylated between NeuN-cells and blood and lineage respectively. Furthermore, comparison of whole brain tissue sections taken from distinct brain regions identified 9 CpG sites that were hypermethylated within cerebellum compared to blood (>80%), lineage (>50%), and 20 additional brain regions/brain-derived cells (>70 %) (Figure 1E). Detailed methods for the selection of all targets, including CpH DNA methylation, can be found within Supplementary Methods. Finally, the fragment length of cfDNA (~166-180pb) indicates that nucleosome-bound DNA is preferentially protected from degradation over non-bound DNA [21]. Therefore increased DNA degradation of blood-cell derived DNA within euchromatic regions of blood cells would be expected during apoptosis by endogenous DNase activity (reviewed in [22]). In an effort reduce background noise (blood cell-derived cfDNA) we prioritized hypermethylated loci within blood euchromatic regions (Supplementary Methods) for targeted assay design (Figure 1F).

**Figure 1.**
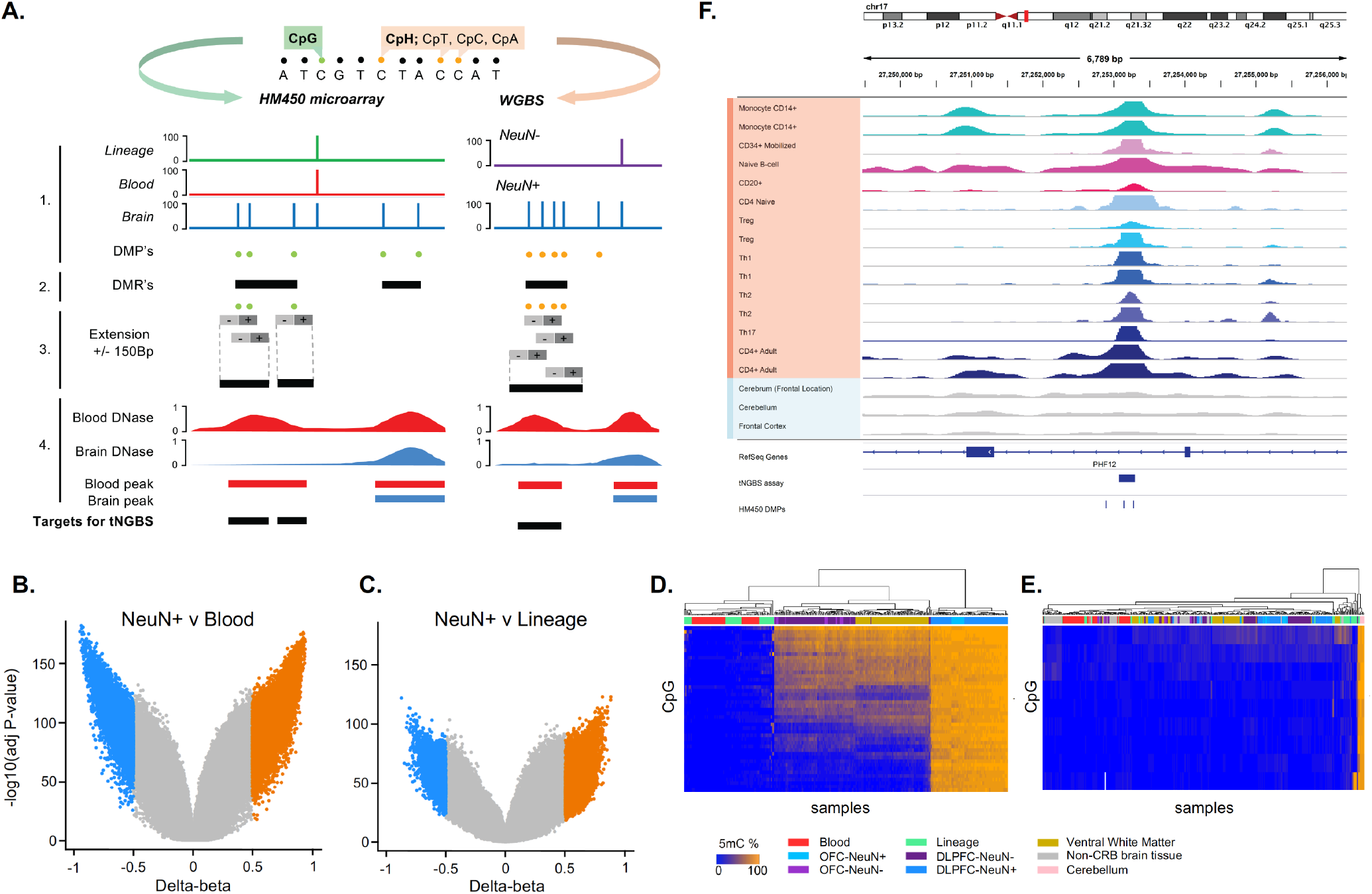
Characterization of brain-specific DNA methylation regions for cfDNA analysis. A. Genomic loci with brain-specific CpG or CpH methylation were characterized using a 4-step approach (Supplementary Material). Step1; DMP’s are identified between brain and blood and brain and lineage tissues (CpG; Illumina HM450K) or DLPFC-NeuN+ and DLPFC-NeuN-(CpH; WGBS), followed by the identification of DMR’s (Step2). Step 3; DMPs intersecting DMRs are extended +/− 150bp. Step 4; Extended DMPs within blood-specific DNAase narrowpeaks represent the target regions for bisulfite assay design. B. Volcano plot of DMP’s identified between NeuN+ cells and Blood. C. Volcano plot of DMP’s identified between NeuN+ cells and Lineage tissues. D. Unsupervised hierarchical clustering of the DNA methylation of 45 CpG found hypermethylated within NeuN+ compared to Blood (>90%) and Lineage (>80%). E. Unsupervised hierarchical clustering of the DNA methylation of 9 CpG found hypermethylated within Cerebellum tissue compared to Blood (>80%), Lineage (>50%) and 20 additional brain regions/brain-derived cells (>70%).

### Validation of targeted bisulfite sequencing assays

Previous groups have shown the utility of bisulfite amplicon sequencing using transposase library construction (BSAS)[23]. We extend from this work and describe protocols for the multiplexing of 35 bisulfite amplicons we term targeted next generation bisulfite sequencing (tNGBS)(Supplementary Materials IV). Bisulfite specific assays were designed to amplify genomic regions found with brain-specific CpG/CpH methylation (Supplementary Materials II). Each assay was validated by analysis of pooled mixtures of CpG methylated and unmethylated DNA standards (CpG assays) and/or pooled mixtures of NeuN+ and PBMC derived DNA (CpG and CpH assays, Supplementary Materials II). The DNA methylation levels of targeted CpG/CpH loci (from discovery) were highly correlated (R^2^>0.8) with the percentage of DNA methylation of the pooled methylation mixtures (Supplementary Figure 4). The tNGBS protocol includes 33 assays targeting genomic regions with brain-specific DNA methylation (Supplementary Methods) in addition to 2 assays targeting spiked in Lambda phage gDNA to control for bisulfite conversion efficiency.

To validate the specificity of the tNGBS assays we performed tNGBS analysis of gDNA from PBMC (N=11), NeuN+ (N=13, DLPFC) and NeuN-(N=7, DLPFC) cells (Figure 2A & B). The analysis revealed highly specific patterns of DLPFC-NeuN+ DNA methylation for assays designed to regions identified wth NeuN+ CpG and CpH methylation. Unsupervised hierarchical clustering of samples resulted in distinct clustering of DLPFC-NeuN+ cells (Figure 2D & E). Unexpectedly we found DLPFC-NeuN+ cells exhibited CpH methylation within regions identified by CpG methylation (Figure 2D), a feature that was also observed in the other 4 assays identified by CpG methylation analysis (Supplementary Figure 5). We performed tNGBS analysis of gDNA from cerebellum and DLPFC whole tissue sections (matched from 3 individuals), Ventral White Matter (N=13), hippocampus (N=2) and PBMC’s (N=11). Unsupervised clustering of tissues revealed distinct clustering of cerebellum tissue (Figure 2F).

**Figure 2.**
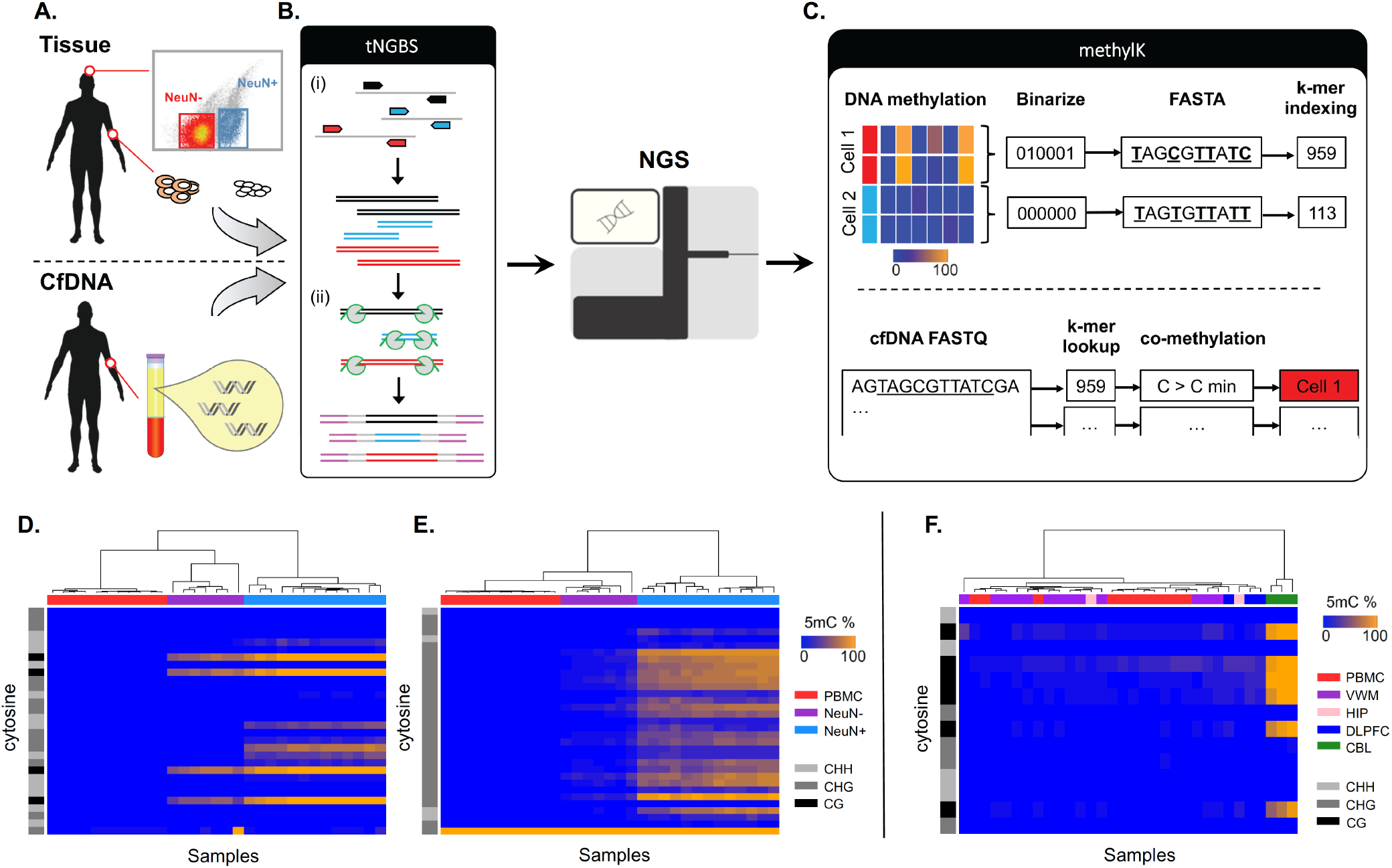
tNGBS-methylK pipeline. A. Validation of brain-specific DNA methylation is performed using genomic DNA (gDNA) from tissue/cells-of-interest (top), for diagnostic application, cell-free DNA is extracted from patient plasma/serum. B. Schematic of targeted next generation bisulfite sequencing (tNGBS) (i) multiplex amplification of bisulfite converted gDNA/ cfDNA produces amplicon pool of regions with brain-specific CpG/CpH methylation. (ii) Tagmentation (Tn5) of the amplicon pool appends 19bp adapters (grey) that are used to append sequencing adapters (purple). The product can then be sequenced using NGS and data analysed by C. methylK pipeline performs two functions; firstly to define DNA methylation k-mers (top) DNA methylation (%) of tissues/ cells-of-interest are binarized (< 50% = 0, > 50% = 1) and converted to FASTA format within the context of surrounding nucleotides (0=T, 1=C). K-mer indexing is then performed on the tissues/cells-of-interest FASTA sequences using Kallisto [20]. Secondly to quantify DNA methylation k-mers (bottom) bisulfite sequencing reads (fastq) from cfDNA samples are used within k-mer lookup of k-mer indexes defined. Correlated-methylation (co-methylation) thresholds based on signal-to-noise ratios are applied to each sequencing read before defining the tissue/ cell-of-origin of the single-molecule. D-E. Unsupervised hierarchical clustering (samples) of tNGBS data shows distinct clustering of DLPFC-NeuN+ cells from PBMC’s and DLPFC-NeuN-cells for example CpG assay (D) and CpH assay (E). F. Unsupervised hierarchical clustering (samples) of tNGBS shows distinct clustering of cerebellum tissue from PBMC’s and other brain regions (VWM, Hippocampus and DLPFC).

### Deconvolution of tissue/cell-of-origin of single molecules using k-mer matching

Researchers have previously used the co-methylated patterns of each single molecule (bisulfite sequencing read) to quantify the tissue-of-origin of cfDNA [10]. However, these methods rely on the computationally intensive alignment of bisulfite sequencing reads to a reference genome, followed by re-analysis of aligned reads for co-methylated cytosines that establish specificity for the tissue-of-origin determination of cfDNA. DNA methylation sites on the single-molecule level are analogous to the unique gene sequences that define transcripts and transcript isoforms. Groups have previously shown that DNA methylation fractions across tissue populations can be used to define co-methylated haplotype blocks for tissue-of-origin detection [24]. Here we establish methods that binarize DNA methylation fractions of tissues/cells-of-interest (methylotype) into FASTA format (Supplementary Figure 6A) that can be used for pseudoalignment of raw bisulfite sequencing reads using the Kallisto framework, termed methylK (https://github.com/zchatt/methylK) (Figure 2C). As proof-of-concept, we performed methylK analysis of PBMC and NeuN+ cells. We found incorrectly pseudoaligned raw bisulfite sequencing reads from either tissue (PBMC pseudoaligned to NeuN+ or NeuN+ pseudoaligned to PBMC) typically displayed lower posterior probabilities (Supplementary Figure 6B & C). Importantly, the fraction of reads pseudoaligned to NeuN+ bisulfite FASTA sequences with posterior probabilities >0.5 was significantly lower (P=3.6 x 10^−5^, Students T-test) in PBMC samples compared to DLPFC-NeuN+ cells (Supplementary Figure 6B). We therefore set a posterior probability threshold of >0.5 for pseudoaligned reads. This threshold retained 76 - 90% of reads originally aligned to NeuN+ assays (bismark) from DLPFC-NeuN+ cells while only 0.18-0.6% of reads were retained from PBMC samples.

The use of correlated DNA methylation has been shown to increase the specificity of cell-of-origin deconvolution of cfDNA [10]. Following pseudoalignment, the number of co-methylated cytosines on each pseudoaligned read for each tissue/cell-of-interest can be counted (Figure 2C & Supplementary Figure 6D). A signal-to-noise ratio (SNR) could then be established by calculation of the proportion of tissue/cell-of-interest pseudoaligned reads with 1:N^th^ methylated cytosines over PBMC samples (Supplementary Figure 6E). We observed high SNRs as high as 66896:1 and applied a 5000:1 SNR threshold for all analysis (Supplementary Figure 6E). Notably, only NeuN+ assays (n=25) and the cerebellum assay passed SNR threshold, none of the DLPFC-NeuN-assays or hippocampus assay passed threshold and were not included within analysis of cfDNA (below).

### Measurements of cfDNA purity within subject samples

Traditionally, bio-banked serum and plasma samples from neurological subjects have not been processed via high speed centrifugation with the anticipation of cfDNA analysis, which has resulted in large background noise (blood cell derived DNA). Typically, cfDNA is ~166-180bp, corresponding to the approximate length of DNA wrapped around one nucleosome core (147bp) plus DNA tails following endogenous DNase activity (~10bp). The length of assays within our tNGBS ranged from 88-245bp. Coverage was highly assay specific across the population of samples analyzed (Supplementary Figure 7A), thus we established a metric, Weighted Reads (WR), to account for assay specific coverage variability (Supplementary Methods III). Generally, PBMC derived gDNA had a higher WR than matched cfDNA samples (WR; Students T-test, P=2.35 x 10^−8^), likely due to degraded cfDNA. Interestingly, we observed a drop-off in the WR of assays >170bp in cfDNA samples compared to PBMC (Supplementary Figure 7B), thus providing a proxy for cfDNA purity (Weighted Read Ratio [WRR], Supplementary Methods III) i.e. the coverage across assays >170bp would be higher in samples with cellular contamination.

We found that the WRR estimates were dependent on sample collection and processing protocol (Supplementary Figure 8). For instance, control samples in which bloods were processed proximal to the time of collection (within 4 hours, Methods) contained a lower WRR than control samples collected in serum tubes (P-value = 0.018). Notably, the lowest WRR were observed in samples collected in specialized cfDNA collection tubes (Qiagen, ccf-DNA). The higher amount of background (non-cfDNA) would be expected to reduce the ability to detect brain-derived cfDNA. Indeed, we found a higher number of DLPFC-NeuN+ cfDNA within samples collected in either ccf-DNA and EDTA tubes processed proximally compared to samples collected in EDTA tubes and processed distally or serum collected samples (P < 0.04, Students T-test).

#### Application of cfDNA analysis by tNGBS-methylK to Acute Blast Exposure, Cognitive Decline and Chronic Neurodegenerative Disease: Army Explosive Entry Personnel (aka Breachers)

CfDNA was isolated from breachers (N=12) on day 1 (prior to training) and on training days 7, 8, and 9. During each training session, operational blast exposure measured by psi-ms was determined by use of mounted pressure monitors (Figure 3A). Single-molecule deconvolution of DLPFC-NeuN+ cfDNA was performed by tNGBS-methylK. We detected DLPFC-NeuN+ cfDNA in eight assays. On day 7 of training the breachers were exposed to significantly higher pressure than other training days (Students T-test, P-value < 1.6 x 10^−13^) as measured by helmet mounted pressure monitors (Figure 3B). Linear regression was performed using the DLPFC-NeuN+ cfDNA fragments/mL of plasma for each subject, contrasting the daily maximum impulse exposure of each individual (psi-ms), representing the acute pressure exposure. Due to the high signal-to-noise ratio, half of residual variances were exactly zero and thus eBayes was unreliable. All DLPFC-NeuN+ assays displayed a positive association (Fold-Change) between daily maximum impulse exposures (Figure 3C), of which 3 assays were significantly associated (p-value < 0.05) with this acute exposure (Figure 3C). We observed an increase of 3 individual DLPFC-NeuN+ assays within an individual following day 7 blast exposure (Figure 3E).

**Figure 3.**
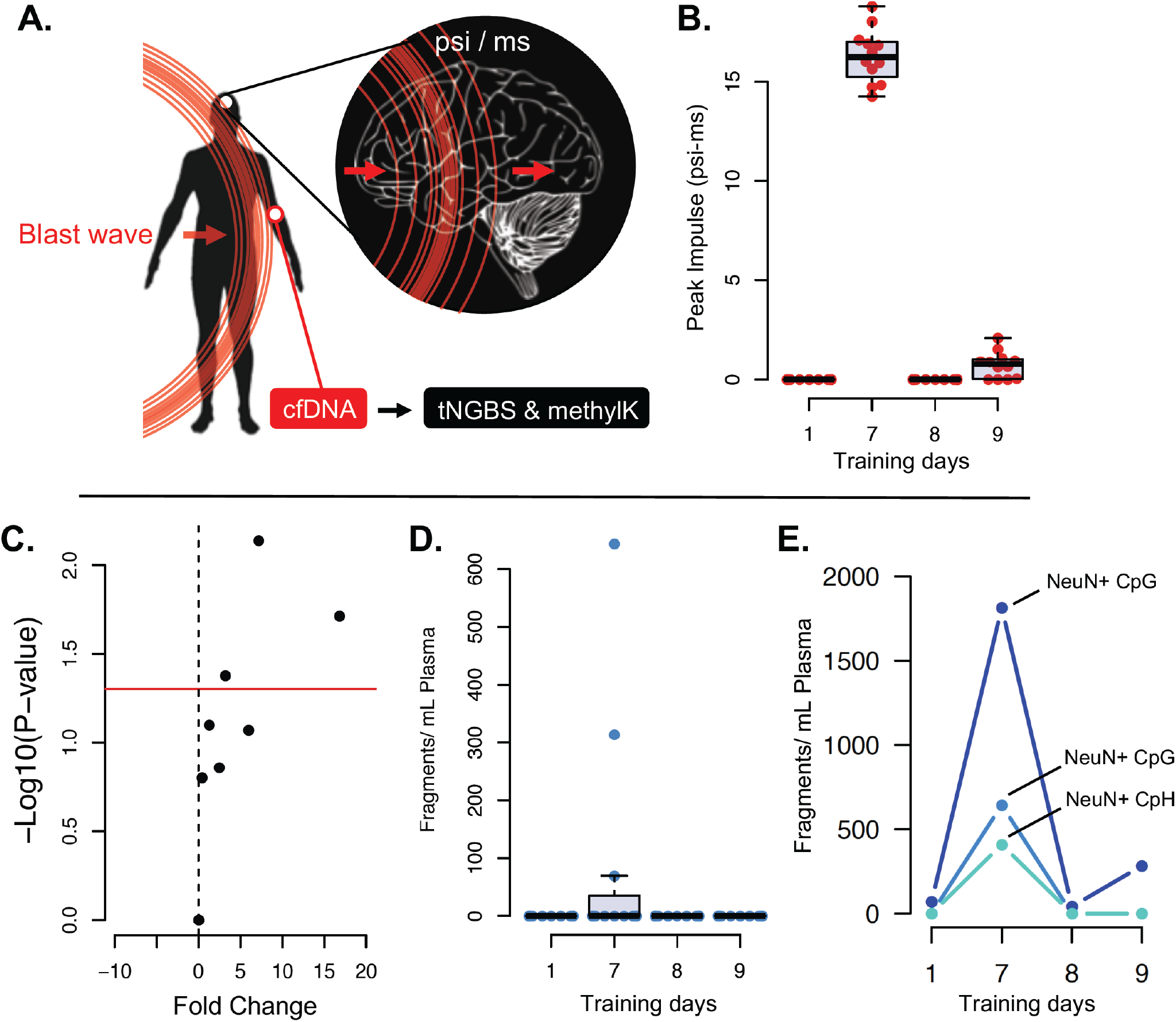
Brain-derived cfDNA detection as a possible indicator of neurotrauma. A. Schematic of study; blast wave exposure of explosive entry personnel was measured (psi-ms) and cfDNA was analysed (tNGBS-methylK) post exposure on days 1,7,8 and 9. B. Blast wave exposure (peak daily impulse) of explosive entry personnel across training in which significantly higher exposure was recorded on day 7. C. DLPFC-NeuN+ cfDNA was detected in the serums of explosive entry personnel and was positively associated with blast wave exposure (Lm) in all 9 assays detected, of which 3 were significant (P-value < 0.05). D. Results from the most significant assay from the Lm (C) targeting CpG methylation of DLPFC-NeuN+ cells in which three individual personnel had detectable DLPFC-NeuN+ cfDNA on day 7 following increased blast exposure. E. Example of an explosive personnel’s DLPFC-NeuN+ cfDNA profile (only assays detectable) over all training days in which a peak in DLPFC-NeuN+ cfDNA was observed following day 7 exposure in all 3 assays.

#### Parkinson’s Disease

We investigated the presence/differences in DLPFC-NeuN+ cfDNA within a chronic neurodegenerative disease, PD, by comparison to non-neurodegenerative “controls”. CfDNA analysis (tNGBS-methylK) was performed in 2 independent cohorts of PD and controls (cohort 1 “Raj” and cohort 2 “Walker”) (Figure 4A). DLPFC-NeuN+ cfDNA was detected by 7 assays in both PD and control subjects within both cohorts. Linear modelling within cohort 1 (PD; n=10, control; n=10) did not reveal any significant differences in the amount of DLPFC-NeuN+ cfDNA between PD and control subjects (Figure 5B). 6/7 assays recorded higher amounts of DLPFC-NeuN+ cfDNA within control subjects, the two assays of highest significance exhibited a fold-change increase of 104 and 675 in controls (P-value=0.15 and 0.3 respectively) (Figure 5C & D). Overall, controls contained higher amounts of DLPFC-NeuN+ cfDNA than PD cases (fold-change=182, DLPFC-NeuN+ cfDNA aggregate).

**Figure 4.**
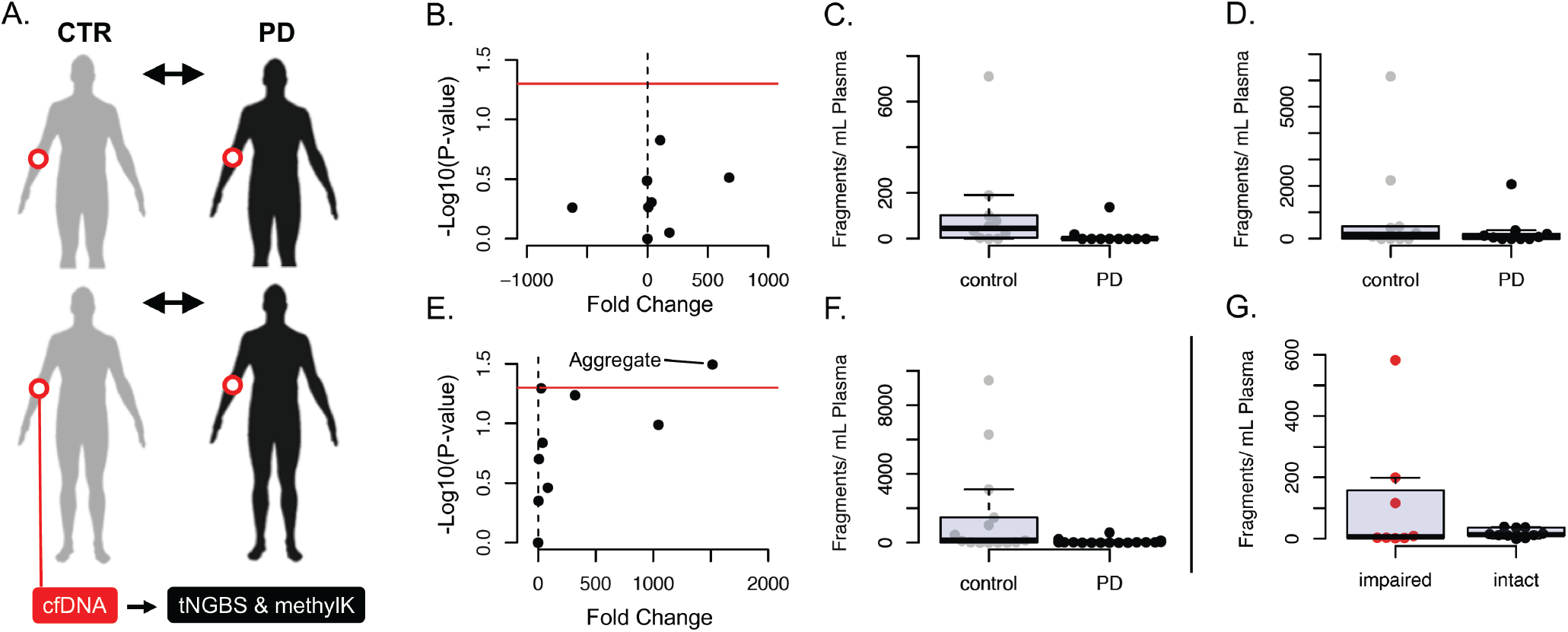
Brain-derived cfDNA detection in Parkinson’s Disease and non-neurodegenerative controls. A. Schematic of study design; Two independent cohorts of PD and non-neurodegenerative controls were compared for brain-derived cfDNA (tNGBS-methylK). B. Volcano plot of P-values and fold change of detected DLPFC-NeuN+ cfDNA between PD and controls in cohort 1 (Raj). C & D Boxplots of detected DLPFC-NeuN+ cfDNA within controls and PD patients for the two DLPFC-NeuN+ assays with highest significance (B). Volcano plot of P-values (Lm) and fold change of detected DLPFC-NeuN+ cfDNA between PD and controls in cohort 2 (Walker). F. Boxplot of aggregate of detected DLPFC-NeuN+ cfDNA within controls and PD patients. G. Boxplot of aggregate of detected DLPFC-NeuN+ cfDNA detected within cohort 2 cognitively impaired PD patients and cognitively intact PD patients.

**Figure 5.**
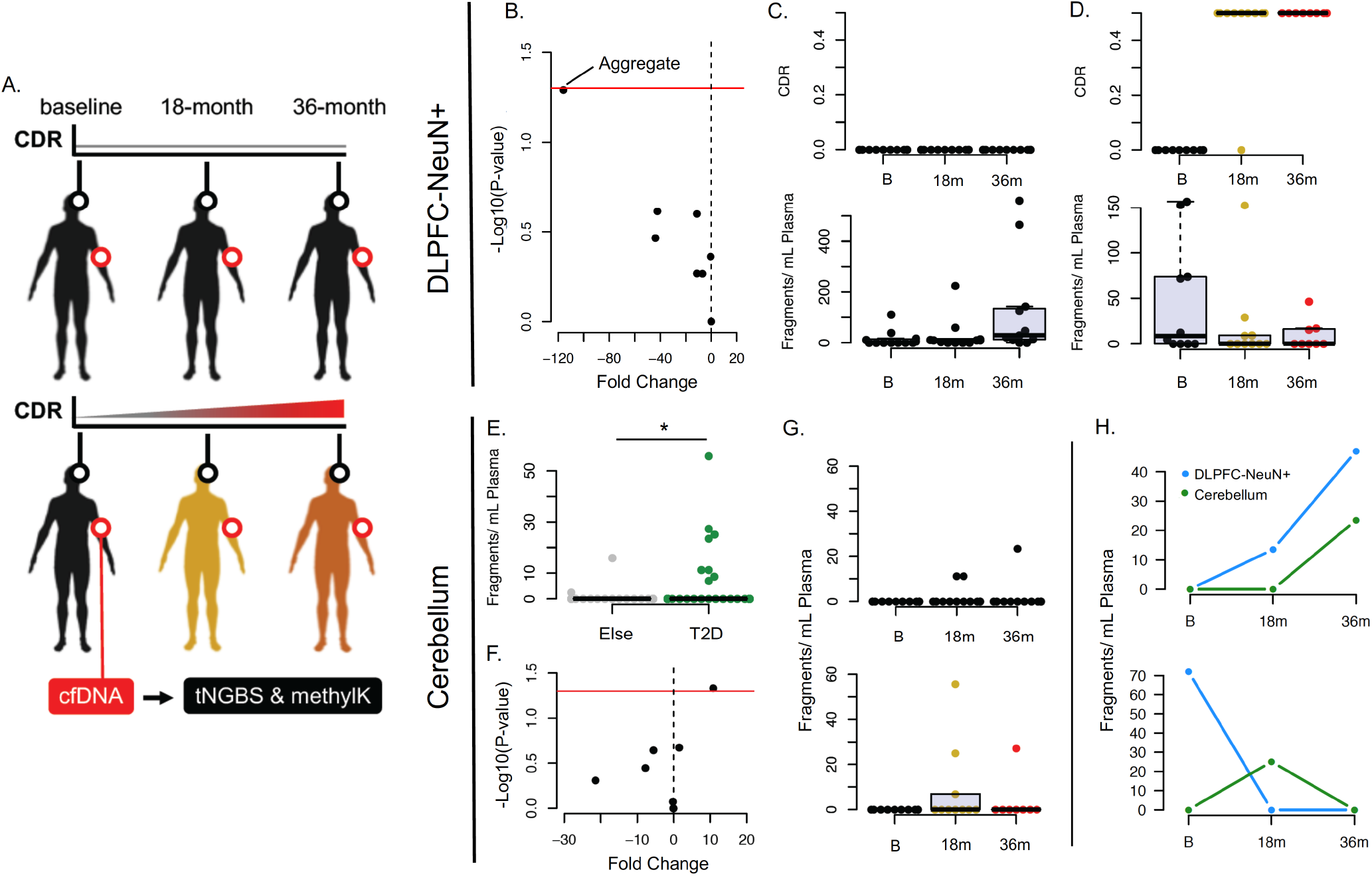
Brain-derived cfDNA detection during cognitive decline in type-2 diabetic cohort. A. Schematic of study design; cognitive assessments were performed in a cohort of type-2 diabetic patients 3 time-points over 36 months and cfDNA was analysed (tNGBS-methylK) at each follow-up. B. Results of linear modelling of DLPFC-NeuN+ cfDNA detected within type-2 diabetic serums and matched Clinical Dementia Rating (CDR) scores. All DLPFC-NeuN+ assays were negatively associated with CDR, an aggregate of all DLPFC-NeuN+ assays was marginally significant (P=0.05). C. CDR scores (top) and aggregate DLPFC-NeuN+ cfDNA results (bottom) from cognitively stable type-2 diabetics (n=12). D. CDR scores (top) and aggregate DLPFC-NeuN+ cfDNA results (bottom) from type-2 diabetics exhibiting cognitive decline over 36 months (n=10). E. Cerebellum cfDNA was detected within the type-2 diabetic cohort (n samples = 64) at significantly higher levels compared to explosive entry personnel and Parkinson’s Disease (n samples = 100) patients’ samples. F. Results of linear modelling of cerebellum cfDNA detected within type-2 diabetic serums and matched CDR scores. G. Cerebellum cfDNA detected within T2D-stable (top) and T2D-decliners (bottom) for the most significant assay detected (F). H. Examples of DLPFC-NeuN+ and Cerebellum cfDNA changes over 36-month follow-ups within T2D-stable (top) and T2D-decliners (bottom). *p-value = 0.03, students t-test.

These results were unexpected as we had hypothesised that subjects with a chronic neurodegenerative disease would exhibit an increase in the amount of brain-derived cfDNA. To explore this further we collected cfDNA from an additional 18 PD subjects and 14 controls within specialised tubes (ccfDNA Qiagen) that reduce background noise (contributed by PBMC gDNA). Again, we observed higher amounts of DLPFC-NeuN+ cfDNA within controls than PD subjects (DLPFC-NeuN+ aggregate fold-change=25.8, P-value=0.03) (Figure 4E & F). Therefore, both PD cohorts exhibited the same directional effects (all but one assay) in which DLPFC-NeuN+ derived cfDNA fragments were higher in controls. In cohort 2, the cognitive capacity of PD subjects were also assessed by a board certified neurologist. Subjects with mild-moderate to marked impairment were classified “impaired” (8/18 subjects), the remaining 10 subjects were classified “intact”. Group-wise comparisons revealed the highest levels of DLPFC-NeuN+ cfDNA within the “impaired” group (mean; impaired=114, intact=18) (Figure 4G), however these results were not significant (P=0.22, Students T-test).

#### Type 2 Diabetes with cognitive impairment

CfDNA was isolated from type 2 diabetics (T2D; N=22) at 3 time-points over a 36-month follow-up, at each time-point cognitive assessments were also performed (CDR) (Figure 5A). At 36 months follow up 10 patients were rated as having mild cognitive impairment (MCI; CDR=0.5) or mild dementia (CDR=1), referred to as “T2D-decliners” (Figure 5D upper). The remaining 12 T2D subjects were cognitively normal (T2D-stable, CDR=0) over the 36 month follow-up (Figure 5C upper). Single-molecule deconvolution of DLPFC-NeuN+ cfDNA was performed by tNGBS-methylK. DLPFC-NeuN+ cfDNA was detected by 7 assays. Linear regression was performed using the DLPFC-NeuN+ cfDNA fragments/mL of plasma for each subject, contrasting CDR score. No assays passed significance (P-value < 0.05), however all assays exhibited the same negative association (fold-change) with CDR (Figure 5B). Notably, an aggregate of all DLPFC-NeuN+ cfDNA detected for each subject was marginally significant (P=0.05) (Figure 5B), of which the highest amount was detected at baseline when T2D-decliners were pre-symptomatic (Figure 5D bottom). Unexpectedly, T2D-stable subjects displayed a group-wise increase in DLPFC-NeuN+ cfDNA at 36 months (Figure 5C bottom). Interestingly, we detected/ observed progressive increases in DLFPC-NeuN+ cfDNA at each follow-up in 50% (6/12) of T2D-stable subjects compared to just 10% (1/10) of T2D-decliner subjects.

Single-molecule deconvolution of cerebellum cfDNA by tNGBS-methylK analysis was performed across all cohorts. Cerebellum cfDNA was detected by 7 assays found to have a signal-to-noise ratio > 5000:1 for cerebellum gDNA compared to PBMC gDNA (original cerebellum assay and 6 NeuN+ assays). The T2D cohort contained significantly higher amounts of cerebellum cfDNA than all other cohorts (P=0.03, Students T-test) (Figure 5E). Linear modelling of cerebellum cfDNA and CDR revealed that one assay (original cerebellum) was significantly associated with CDR (P=0.04) (Figure 5F). The highest amounts of cerebellum cfDNA were observed at the 18-month follow-up of the T2D-decliners, the time-point in which cognitive decliners first reported CDR scores >0.5 (Figure 5G). The detection of cerebellum cfDNA within 2/3 T2D-stable (3 serum samples) coincided with the detection of DLPFC-NeuN+ cfDNA (Figure 5H [top]), similarly the detection of cerebellum cfDNA within 2/3 T2D-decliners (4 serum samples) coincided with the detection of DLPFC-NeuN+ cfDNA. The detection of cerebellum cfDNA was preceded by DLPFC-NeuN+ cfDNA in the remaining T2D-stable and decliner subjects (Figure 5H [bottom]). Therefore, the detection of cerebellum cfDNA did not precede the detection of DLPFC-NeuN+ cfDNA.

## DISCUSSION

### Characterization and validation of brain-specific CpG/ CpH methylation

CpH methylation is obtained postnatally [11] and contributes largely to cell specification of neuronal subtypes [25]. However, it has not been established if these DNA methylation signatures are common across the population and whether they are stable and maintained throughout advanced aging, when neurodegenerative disease is more likely to occur. Here we validate CpH regional methylation at single-base resolution, representing, to our knowledge, the largest cohort of single-base resolution validation of human neuronal CpH methylation signatures and find highly consistent patterns of the CpH methylation distinct to all DLPFC-NeuN+ cells analysed. We also validate NeuN+ CpG methylated loci detected by genome-scale CpG microarray and observe distinct clustering of DLPFC-NeuN+ cells that was, unexpectedly, driven by CpH methylation within the same genomic loci. Considering that CpG methylation was shared between DLPFC-NeuN+ and NeuN-cells it is possible that local DNA methyltransferase activity common to DLPFC cell-types may lead to spontaneous DNA methyltransferase activity outside of the CpG context within post-mitotic neurons.

We were able to validate the first DNA methylation assay with brain-region specificity to the cerebellum using CpG loci identified from CpG methylation microarray. Considering that CpH methylation occurs ~20x more frequently than CpG methylation in human neurons [11], it is highly likely that CpH methylation will be more discriminatory of DNA derived from brain regions. Thus WGBS of neurons from brain regions implicated in specific neurodegenerative diseases (e.g. the substantia nigra in PD) has the high potential to lead to disease-specific biomarkers. It is worth noting that CpH methylation standards are not currently available. To initially screen for efficient bisulfite assays designed to genomic regions of CpH methylation we serially pooled DLPFC-NeuN+ gDNA with PBMC gDNA from a single subject. Considering the effectiveness of CpH methylation in characterising neuron derived DNA, and the ability of CpH methylation to discern neuronal subtypes [1], the synthesis of a set of genome-wide CpH methylation standards will be essential in expanding cfDNA analytics to discern indications of neurotrauma and track prodromal and long-term neurodegeneration of specific cortical-regions/ layers.

Genome-wide chromatin states are highly cell-type-specific [26], of which a large number have been characterized by exploiting the properties of enzymes that preferentially digest open chromatin (i.e. DNase [27] and transposase [28]) followed by Next Generation Sequencing (NGS). Such profiles are analogous to the endogenous DNase mediated DNA degradation that occurs during apoptosis (reviewed in [22]). Blood-cell-derived cfDNA contributes to the large majority of cfDNA, representing an enormous background signal against which to identify rare cfDNA derived from other cell-types such as neurons. Here we present methodology for the inclusion of cell-specific chromatin profiles in the identification of DNA methylation regions for cfDNA analysis and believe this strategy will benefit assay design in the fields of oncology and transplantation monitoring and may help expand cfDNA technology to disease monitoring in rare disease with limited cfDNA contribution. In addition, blood samples collected from subjects suffering from possible neurotrauma/degenerative disease have traditionally not been collected/ processed using methods that limit background noise (blood derived cfDNA). We provide a method (WRR) that can establish the amount of non-cfDNA background that we hope will aid other researchers in determining the purity of cfDNA within their plasma/ serum biobanks.

### Technological developments for cfDNA methylation analysis

Here we present multiplex bisulfite NGS protocols (tNGBS) that are capable of simultaneous analysis of 35 genomic regions. The benefit of DNA methylation biomarkers of cell-type is that numerous sites can be used to identify the same cell type, improving assay specificity through internal validation. Considering the diversity of cell-types within the human body, we anticipate that cfDNA methylation analysis will be expanded to incorporate many genomic-sites of interest of various cell types. This will pose a strain on computational methods that rely on the alignment of bisulfite sequencing reads to a reference genome. Thus we present new analytical methods (methylK) for the analysis of cfDNA methylation. Binarization and k-mer based lookup represent an ultra-fast method of single-molecule deconvolution that will scale with the increase in targeted loci anticipated. Further, methylK analysis should have a variety of applications outside of cfDNA methylation analysis, i.e. analysis of any tissue with a variety of cell-types.

### Application of cfDNA methylation analysis in possible neurotrauma and neurodegenerative disease

Breachers: Injuries from exposure to explosive blasts rose dramatically during the Iraq/Afghanistan Wars, motivating investigations of blast-related neurotrauma. In this effort, we have undertaken human studies involving “breachers”, military and law enforcement personnel who are exposed to repeated blast as part of their occupational duty. Breachers are typically in close proximity to controlled, low-level blast during explosive breaching operations and training, being exposed to blast overpressure waves. Career breachers (with upwards of hundreds of blast exposures) have reported a range of physical, emotional, and cognitive symptoms, including headache, sleep issues, anxiety, and lower cognitive performance, akin to some symptoms of mild TBI. To our knowledge, we report the first evidence of DLPFC-NeuN+ cfDNA following low-level blast exposure within a longitudinal operational breaching training. During the course of the training, blast overpressure was measured using mounted pressure monitors. It should be noted that the breacher personnel did not report clinical symptoms associated with mild-TBI following exposure. Brain-derived cfDNA biomarkers represent a new class of biomarker for indication of possible acute brain injury and warrant validation in additional sub-clinical cohorts. Further, this technology offers a framework to advance the application of brain-derived cfDNA tNGBS to sports-related concussion.

Parkinson’s Disease: There are currently no validated biofluid biomarkers in clinical use for PD, although several peripheral blood markers have been proposed as candidates, such as α-synuclein, DJ-1, uric acid, and epidermal growth factor (EGF) (reviewed in [29] and [30]). There is a critical need for the identification of biomarkers of neurodegeneration that are economical and can thus be administered to pre-symptomatic patients, e.g. those with familial risk factors, and to track disease progression in clinical trials. Here we perform, to our knowledge, the first analysis of brain-derived cfDNA in PD. Our data, replicated in 2 distinct PD cohorts, indicate that DLPFC-NeuN+ cfDNA in PD subjects is lower than in age-matched non-neurodegenerative controls.

A number of explanations could explain these observations. Firstly, the numbers of dopaminergic neurons lost prior to clinical presentation has been estimated to be from 48-91% [31]. Our PD subjects may have already experienced a significant amount of neurodegeneration, thus the number of residual degenerating neurons contributing to brain-derived cfDNA would be limited. Another potential explanation is that the primary neurons undergoing degeneration in PD (dopaminergic neurons of the substantia nigra pars compacta) do not share DNA methylation profiles with DLPFC-NeuN+ for which our assays were established. Considering the role of the prefrontal cortex in executive functions and the selective degeneration of the prefrontal cortex in AD [32], the detection of DLPFC-NeuN+ cfDNA may be more specifically associated with prefrontal cortex degeneration and cognitive dysfunction. In line with this theory, we observe a trend of higher DLPFC-NeuN+ cfDNA in PD subjects with mild-moderate cognitive impairment compared to cognitively intact subjects. Further studies are required to investigate differential markers of various cortical regions, as certain regions of cortex are known to also be affected in PD [33].

Type 2 diabetes: Patients with type 2 diabetes have an increased risk of developing mild cognitive impairment, vascular dementia and AD [34, 35]. The etiological mechanisms are multifactorial, however hyperglycemia and vascular disease are likely to be involved (reviewed in [36]). Considering the heightened risk, it is imperative to have an economical test of neurological damage that can be applied at a population scale to identify patients within the pre-symptomatic stages of cognitive decline. Here we perform the first analysis, to our knowledge, of brain-derived cfDNA in type 2 diabetics with cognitive impairment. Through intra-individual longitudinal cognitive assessments, we were able to assess patients undergoing cognitive decline. Importantly the study design enabled us to collect and analyze pre-symptomatic samples prior to the onset of significant cognitive impairment. We report a negative association of DLPFC-NeuN+ cfDNA with CDR that is indicative of an active period of neurological damage prior to cognitive decline. In T2D-decliners the highest amount of DLPFC-NeuN+ cfDNA was detected 18-months prior to cognitive decline. We also detected DLPFC-NeuN+ cfDNA within the T2D-stable subjects; interestingly the highest amounts detected were at the last follow-up visit. Future follow-up evaluations of this cohort will be important to establish if these T2D-stable subjects exhibiting detectable DLPFC-NeuN+ become cognitively impaired.

We also report the first evidence of cerebellum-derived cfDNA. The detection of cerebellum cfDNA was almost exclusively found within type 2 diabetic cohort and was exclusively detected at or after the detection of DLPFC-NeuN+ cfDNA. Structural imaging studies have shown that cerebellar grey matter atrophy is a feature in the progression of AD, however functional neuroimaging have shown that there is an increase in cerebellar activation in MCI and AD which has been suggested to represent a compensatory mechanism (reviewed in [37]). It is therefore plausible that early detection DLPFC-NeuN+ cfDNA and subsequent detection of cerebellum cfDNA represents a quantifiable peripheral biomarker reflecting cognitive reserve. This could represent important metrics for staging of subjects within the disease course. This requires validation within larger cohorts and future efforts should also focus on the development of additional cerebellum cfDNA assays.

### Looking ahead

Brain-derived cfDNA represents a new class of peripheral biomarkers that can characterize the cell-type and brain region affected by neurological damage. While this is a proof-of-concept study, the pipeline we establish has great potential to be extended to various neuronal cell types and brain regions. One limitation of the discovery phase (at the time we performed this study) was the availability of WGBS data for various cell types of the human brain, particularly within regions affected in common neurodegenerative diseases. Characterization of DNA methylation within tissues/cell-types that are distinctly affected in different neurodegenerative diseases (i.e. substantia nigra in PD) holds the potential for establishing disease-specific assays. However, many questions remain to be answered with respect to each neurological condition. At what point during disease progression is brain-derived cfDNA detectable? Are there peak times during disease development? Is there any diurnal variability? Is cfDNA from different brain regions detectable at different disease stages? Can these markers be used in therapeutic trials to identify appropriate times to administer treatments or to monitor disease progression? CfDNA analysis is a blood-based method that can be implemented in large populations and with great frequency which will enable epidemiological analysis; do particular genetics, dietary, lifestyle activities etc. lead to an increase in brain-derived cfDNA? Is brain-derived cfDNA a risk factor for disease? Unfortunately, there are more questions than answers at the moment, but the potential is vast.

## METHODS

### Subjects and Sample Processing

Postmortem specimens: Dorsolateral prefrontal cortex (DLPFC) and ventral white matter (VWM) were dissected from cases diagnosed with either schizophrenia (n=24), major depressive disorder (n=24), or control subjects with no known neuropsychiatric disorder (n=24). No subjects had been diagnosed with any neurodegenerative disease at time of death. Subjects had a broad age range (22-72 years) with a mean age of 50 years (subject details in Supplementary Table 1).

Blood-derived samples and processing: Breacher Subjects; In collaboration with the Walter Reed Army Institute of Research, to investigate the effects of repeated blast wave exposure in explosive entry personnel (Breachers), serum samples were taken from 12 breachers following training days 1,7,8 and 9. All samples were from male subjects with an average age of 29.67 yrs (SD 3.87 yrs) (Supplementary Table 10). Briefly, serum was separated by centrifugation at 3000g for 15min at RT and was clarified by additional centrifugation at 10,000g for 10min; 4C. The extracted serum was stored at −80C until cfDNA extraction and tNGBS processing (Supplementary Protocol).

Parkinson’s Disease Subjects;_Cohort 1: Whole blood was collected within 10mL EDTA tubes and processed within 3 hours of collection. Blood was centrifuged at 1700 rpm for 10 minutes and plasma was pipetted 1mL aliquots at −80C. Additional subject details can be found in Supplementary Table 9.

Cohort 2: Whole blood was collected from a separate cohort of 18 subjects within PAXgene Blood ccfDNA Tubes (Qiagen). Blood plasma was separated by centrifugation for 15 minutes at 1900g and clarified with a second centrifugation for 10 minutes at 1900 x g. Subjects cognitive impairment was assessed by clinical evaluation by board-certified neurologist (RHW). Additional subject details can be found in Supplementary Table 6.

Control Subjects; Whole blood was collected from 14 subjects with no known neurological or neurodegenerative disorders as part of an ongoing longitudinal study of suicidal behavior. Blood samples were collected and processed as described for PD Cohort 2. Additional subject details can be found in Supplementary Table 7.

Subjects with Type 2 Diabetes with Cognitive Decline; As part of the Israel Diabetes and Cognitive Decline (IDCD) study [12], whole blood was collected from subjects within serum tubes and stored on ice for a maximum of 6 hours before laboratory processing. The blood tubes were centrifuged (10 minutes, 1,500 rpm) at room temperature. Serum was then pipetted from top and stored at −80C. All subjects were diagnosed with Type 2 diabetes. Cognitive assessments were performed using the Clinical Dementia Rating (CDR) [13] at baseline and each follow-up to measure cognitive decline. Bloods and CDR were assessed during the same visit. Two groups of participants were selected: Decliners (N=10), i.e. participants whose CDR declined from a status of no dementia (CDR=0) to a status of questionable dementia (CDR >= 0.5) or frank dementia (CDR>1) at two consecutive follow-ups (18 and 36-months). The second group was of control subjects in which CDR scores = 0 at all three follow ups follow-up. Additional subject details can be found in Supplementary Table 8.

### Fluorescent activated cell-sorting (FACS)

Neuronal nuclei were isolated from the DLPFC dissections using methods previously described [14]. Briefly, frozen sections of the DLPFC (~100mg) from each subject (N=72) were homogenized on ice, cells were lysed using a cell lysis buffer, and nuclei were isolated by highspeed centrifugation through a sucrose buffer. Nuclei were immunostained using the Alexa-Fluor conjugated anti-NeuN antibody (Abcam, ab190195) and isolated by FACS (BD FACS-Aria).

### DNA methylation microarray profiling

DNA extracted from all 216 (72 NeuN+ DLPFC, 72 NeuN-DLPFC and 72 Ventral White Matter [VWM]) samples was subjected to genome-wide DNA methylation interrogation by the Illumina HM450 DNA methylation microarray, of which 192 passed QC (described below). The analyses were performed using R Language 3.03[15], an environment for statistical computing, and Bioconductor 2.13[16]. Raw data files (.idat) were processed by minfi package [17]. All samples displayed a mean probe-wise detection call p-value for the 485512 array probes < 0.0005. Beta-values (logit transformed M-values) were used for DNA methylation reporting. Sample removal:samples were removed that did not match phenotypic sex and methylation-based sex calling using the getSex function of minfi. Samples with non-matching genotypes (HM450 SNP interrogating probes) between tissues of the same individual were also removed; 192 samples were retained following sample removal (Supplementary Table 1).

### Publicly Available Data for Methylated cfDNA Target Identification

DNA methylation microarray data: We obtained 63 additional publicly available HM450 datasets from different brain regions generated by the BrainSpan consortium and 24 HM450 NeuN+/− derived from the occipital frontal cortex (OFC) (GSE50798) (Supplementary Table 2), resulting in a joint collection of 279 DNA methylation microarray profiles from the human brain. Publicly available HM450 data from peripheral blood (GSE41169 & GSE32148), various primary and cultured peripheral tissues (GSE29290, GSE20945 & ENCODE) derived from the 3 germ layers were obtained, termed “Lineage” (Supplementary Table 2).

DNAse hypersensitivity data; DNAse narrow peak datasets produced from brain and blood samples by the ENCODE consortium to identify brain-specific euchromatic regions are described in Supplementary Table 3.

Whole Genome Bisulfite Sequencing (WGBS) data; WGBS data from FACS sorted (NeuN+ and NeuN-) DLPFC from 2 adults (1 male, 1 female) were generated by Lister et al (GSE47966).

### Identification of brain region and cell specific methylated cfDNA targets

cfDNA targets derived from CpG methylation microarray data: We devised a generalizable 4-step pipeline (Figure 1A) for the identification of loci and /or genomic regions displaying brain region or brain cell specific DNA methylation - the analysis was performed for each brain region or cell type of interest separately. Differential Methylated Position (DMP) analyses were performed using LIMMA [18] and Differential Methylated Region (DMR) analyses within brain regions/cells and contrasting non-brain tissue and cell types were performed using bumphunter algorithm [19]. The brain region/cell specific DMRs were further screened and filtered using overlapping DNase hypersensitivity datasets, contrasting chromatin accessibility peaks between brain vs. blood.

CfDNA targets derived from CpH methylation from NeuN+/NeuN-Whole Genome Bisulfite Sequencing data: We utilized adult WGBS data produced from the prefrontal cortex (1 male, 1 female) that was FACS separated into NeuN+ and NeuN-as described in Lister et al (GSE47966). CpH cytosines with >4X coverage were retained and hyper/hypomethylated cytosines were defined as 100/0% methylated respectively. We identified cell-specific CpH hypermethylation sites by comparison of hyper-hypomethylated CpH cytosines between each cell-type. The number of hypermethylated CpH sites was then calculated within +/-50bp of each cell-specific hypermethylated CpH which defined the density of CpH hypermethylation. Genomic regions were prioritized for targeted assay design based on the density of hypermethylated CpH loci.

### Multiplex Bisulfite Sequencing cfDNA Target Regions Assay Design and Testing is described in detail in Supplementary Materials II

#### Targeted Next Generation Bisulfite Sequencing (tNGBS)

CfDNA extraction, Bisulfite conversion, PCR and NGS library construction necessary for tNGBS are described in detail in the Supplementary Materials II. As reference for the tNGBS analyses, postmortem brain specimens were included and sequenced. Specifically, matched DLPFC (n=3) and cerebellum (n=3) tissues were dissected from 3 subjects (Supplementary Table 5). DLPFC NeuN-(n=6), NeuN+ (n=12) and VWM (n=11) also interrogated by HM450 microarray were also analyzed (Supplementary Table 1). Additionally, details of the tNGBS analyzed samples and sequencing are provided in the Supplementary Tables 4-10. Libraries were denatured following Illumina protocols and sequenced on either the Illumina MiSeq using 2 x 26bp PE sequencing or Illumina HiSeq2500 using 2 x 50bp PE sequencing. Additional details can be found within Supplementary Table 4.

Processing of tNGBS data: Paired-end 50bp reads were trimmed to 26bp using fastqutils truncate function for combined analysis with paired-end 26bp reads. Fastq files were processed using cutadapt to trim poor-quality reads (< Q30). A targeted bisulfite reference genome was generated using bismark command bismark_genome_preparation for the fasta file that includes all 35 assay targets and Lambda genome. Trimmed paired-end reads were then mapped to the targeted bisulfite reference genome using bismark. Stacked DNA methylation calls and coverage were obtained using bismark_methylation_extractor (-p -comprehensive --merge_non_CpG).

Quality Control tNGBS: The mean DNA methylation percentage of all cytosines within Lambda assays were used to determine the conversion efficiency of each bisulfite conversion reaction. We found that 24 of the 254 samples analyzed in this study had a bisulfite conversion efficiency < 98%, including 1 VWM, 2 DLPFC NeuN+ and 3 PBMC and 18 subject cfDNA samples. We removed the whole tissue samples with <98% conversion efficiency from further analysis and annotated the subject cfDNA samples.

K-mer analysis of raw bisulfite sequencing reads: Cell-type specific DNA methylation patterns are analogous to gene sequences that delineate transcripts; gene sequence substrings (k-mers) can delineate between transcripts, similarly bisulfite sequencing k-mers are unique between DNA fragments exhibiting cell-specific DNA methylation patterns within a pooled mixture of DNA. Recently Bray and colleagues [20] described Kallisto, an ultra-fast method that matches k-mers in raw sequencing reads to transcript specific k-mers using a hash table. We establish wrapper scripts to make Kallisto applicable to bisulfite sequencing cell-of-origin detection, outlined within Figure 4. These scripts are available at https://github.com/zchatt/methylK. Briefly, the DNA methylation of all cytosines within genomic regions that exhibit cell-specific DNA methylation were binarized (methylated >0.5, unmethylated <0.5) using the cytosine DNA methylation fractions derived from analysis of the tissue/cell-of-interest, referred to as methylotypes. Methylotypes, stored in .vcf format, were sorted (Picard) and converted to FASTA format using gatk -T FastaAlternateReferenceMaker. The FASTA files were then indexed using Kallisto software [18]. Pseudoalignment of raw FASTQ reads were then performed using Kallisto [18]. Posterior probability thresholds (>0.5) were applied to pseudoaliogned reads to reduce false-positive pseudoalignment. Additionally, methylated cytosines were counted within each pseudoaligned read. Signal-to-noise thresholds were calculated for pseudoaligned reads with 1 to 26 (read length) co-methylation events by comparing the fraction of pseudoaligned reads with co-methylation events between brain tissue/cell and PBMC e.g. NeuN+ and PBMC. All pseduoaligned reads used in this study were found to have signal-to-noise thresholds >5000:1. Pseudoalignment of raw bisulfite sequencing reads from cfDNA defines the cell/tissue of origin. To normalize the data, pseudoaligned reads from cfDNA reads were divided by the cfDNA concentration (the ratio of total ng cfDNA extracted to total mL plasma or serum used for extraction).

## AUTHOR CONTRIBUTIONS

Z.C and F.H conceived the study and coordinated experiments. A.D performed post-mortem brain tissue dissections. Z.C, N.M, S.C performed nuclei isolation and fluorescent activated nuclei sorting. Z.C and Y.G performed bioinformatic and statistical analysis. Z.C performed tNGBS analysis, conceived and programmed the methylK software. W.C and G.K recruited and collected breacher subjects Materials & Correspondence for this study. M.B recruited and collected type 2 diabetic patients for this study. R.W and T.R recruited and collected PD patients for the study. R.W performed cognitive assessments of the PD cohort 2. Z.C and F.H wrote the manuscript, with contributions from all authors. All authors read and approved the final manuscript.

## ACKNOWLEDGEMENTS

We thank Liying Yan (EpigenDX) and her team for their help in assay design and synthesis. This work was supported in part through the computational resources and staff expertise provided by Scientific Computing at the Icahn School of Medicine at Mount Sinai. This study was supported by a postdoctoral fellowship (Z.C) of the NIDA T32 postdoctoral training program in Interdisciplinary Training in Drug Abuse Research and VA Merits I01RX001705 and I01CX001395. Sample collections (G.K) were supported by Broad Agency Announcement Award No. W81XWH-16-2-0001. Material has been reviewed by the Walter Reed Army Institute of Research. There is no objection to its presentation and/or publication. The opinions or assertions contained herein are the private views of the authors and are not to be construed as official, or as reflecting true views of the Department of the Army or the Department of Defense. The investigators have adhered to the policies for protection of human subjects as prescribed in AR 70-25. This research was supported in part by an appointment to the Research Participation Program at the Walter Reed Army Institute of Research administered by the Oak Ridge Institute for Science and Education through an interagency agreement between the U.S. Department of Energy and USAMRMC.

## MATERIALS AND CORRESPONDENCE

All correspondence and material requests should be addressed to Z.C and F.H

## ABBREVIATIONS

(cfDNA): Cell-Free DNA
(DLPFC): Dorsolateral Prefrontal Cortex
(HM450): Illumina Infinium Human Methylation Microarray 450K platform
(WGBS): Whole Genome Bisulfite Sequencing
(DMR): Differentially Methylated Region
(DMP): Differentially Methylated Positions
(tNGBS): Targeted Next Generation Bisulfite Sequencing
(TBI): Traumatic Brain Injury
(PD): Parkinson’s Disease
(CDR): Clinical Dementia Rating
(NGS): Next Generation Sequencing

