## Supplementary Material for "Brain-derived circulating cell-free DNA defines the brain region and cell specific origins associated with neuronal atrophy"

**Appendix**

I. Selection of genomic regions exhibiting brain-specific DNA methylation
II. Multiplex bisulfite sequencing assay design and testing.
III. Measurements of cfDNA purity using Weighted Fraction Ratio's (WFR)
IV. Laboratory protocols for targeted next generation bisulfite sequencing (tNGBS) of cfDNA

#### I. Selection of genomic regions exhibiting brain-specific DNA methylation

*Strategy for characterization of genomic region/ cytosine loci selection for cfDNA analysis*: 4 Step pipeline (Figure 1A);

1. Identifying brain-derived DMPs as compared to blood and other potential tissues contributing to the cfDNA content in plasma.
   i) DMP analysis was performed between brain – blood. Significant DMPs with FDR adjusted $P < 0.05$ & differential methylation > 80% were retained.
   ii) DMP analysis was performed between brain – Lineage. Significant DMPs with FDR adjusted $P < 0.05$ were retained and ranked according to magnitude of methylation difference.
2. Delineating brain specific DMRs to characterize brain region specific methylation compared to blood and other potential tissues contributing cfDNA to plasma.
   i) DMR analysis was performed between brain – blood. Significant DMRs with fwer< 0.05 were retained.
   ii) DMR analysis was performed between brain – Lineage. Significant DMRs with fwer < 0.05 were retained.
   iii) An intersection of DMRs between 2i and 2ii was performed using GenomicRanges and intersected loci were retained.
3. An intersection was performed using GenomicRanges between BOI (brain-of–origin) retained DMP and DMRs to refine DMPs with regional DNA methylation specific to brain. Intersected DMPs were retained within GenomicRanges object after which 150bp +/- bp were added up/ downstream (GenomicRanges150).
4. Defining BOI DMRs (GenomicRanges150) within euchromatin of blood and heterochromatin of brain.
   i) The GenomicRanges150 object was intersected with Blood DNase hypersensitivity narrow peaks (Supplementary Table 3) using GenomicRanges. Intersections were ranked by 1) the number of blood DNase datasets that intersect the combined narrow peak and 2) the enrichment means of blood DNase datasets contributing to each narrow peak. The top $10^{th}$ percentile of peaks by dataset counts and top $10^{th}$ percentile by dataset enrichment were retained.
   ii) The retained DMRs were intersected Brain DNase hypersensitivity narrow peaks (Supplementary Table 3) and binned into intersecting and non-intersecting.
   iii) The CG density of surrounding the DMP within the intersecting and non-intersecting bins was calculated and the bins were ranked from highest density to lowest and selected for targeted sequencing design.

Detailed description of the identification of the final 33 assays included within the multiplex tNGBS.

Assays Overview:

**CpG**

- 4 CpG assays for NeuN- that were identified in NeuN- analysis of CpG (CpG-glia)
- 1 CpG assay with brain-region specificity for the Cerebellum (CpG-Cerebellum)
- 1 CpG assay with brain-region specificity for hippocampus (CpG-region3)
- 4 CpG assays hypermethylated in DLPFC NeuN+ (hyperCpG-neuron)
- 4 CpG assays for DLPFC NeuN+ within DNase hypersensitivity regions (dnaseCpG-neuron)

**CpH**

- 5 CpH assays common to DLPFC NeuN+ and NeuN- (CpH-neuronglia)
- 3 CpH assays for DLPFC NeuN+ with high density of hypermethylated CpH density (highdensityCpH-neuron)
- 8 CpH assays for DLPFC NeuN+ with mid density of hypermethylated CpH density NeuN+ (middensityCpH-neuron)
- 3 CpH assays for DLPFC NeuN+ within DNase hypersensitivity regions (dnaseCpH-neuron)

CpG-glia: To define NeuN- DNA methylation for the detection in NeuN- cfDNA we contrast NeuN- cells (n=78; DLPFC and OFC) genome-wide DNA methylation profiles (Illumina, HM450) to PBMC and Lineage genome-wide DNA methylation profiles (Illumina, HM450) using LIMMA separately. We identified 13 CpG sites (distinct regions) that exhibited FDR < 0.05 in both analysis and > 0.8 delta-beta between NeuN- and Blood and > 0.5 delta-beta between NeuN- and Lineage tissues. Four tNGBS assays were designed and tested from the 13 sites (Supplementary Figure 1).

CpG-Cerebellum: To define Cerebellum DNA methylation for the detection in Cerebellum cfDNA we contrast Cerebellum (5) genome-wide DNA methylation profiles (Illumina, HM450) to PBMC and Lineage genome-wide DNA methylation profiles (Illumina, HM450) using LIMMA separately. We identified 716 CpG sites that exhibited FDR < 0.05 in both analysis and > 0.8 delta-beta between Cerebellum. We then contrast Cerebellum genome-wide DNA methylation profiles (Illumina, HM450) to all other brain derived genome-wide DNA methylation profiles (274) (Illumina, HM450) and identified 9 CpG sites that exhibited FDR < 0.05 and delta-beta of > 0.7 as well as a delta-beta > 0.5 between Cerebellum and Lineage (previous analysis). One tNGBS assay was designed and tested (Figure 2F) from the 9 sites (Figure E).

CpG-region3: To define DNA methylation specific to the Hippocampus (HIP) and inferolateral temporal cortex (ITC) we contrast genome-wide DNA methylation of the HIP (3) and ITC (6) to PBMC and Lineage tissues using separately using LIMMA. We identified 67 CpG sites that exhibited FDR < 0.05 and delta-beta > 0.6 between HIP/ITC and PBMC. We then performed analysis between (LIMMA) between HIP/ITC genome-wide DNA methylation and genome-wide DNA methylation profiles of brain tissues derived from the Frontal Cortex (IPC +STC +MFC + VFC +DFC +OFC). We identified 5 CpG sites with FDR < 0.05 and delta-beta > 0.1 between HIP/ ITC and Frontal Cortex and delta-beta > 0.5 between HIP/ITC and Lineage tissues.

HyperCpG-neuron: To define NeuN+ DNA methylation for the detection in NeuN+ cfDNA we contrast NeuN+ cells (n=71; DLPFC and OFC) genome-wide DNA methylation profiles (Illumina, HM450) to PBMC and Lineage genome-wide DNA methylation profiles (Illumina, HM450) using LIMMA separately. We identified 45 CpG sites that exhibited FDR < 0.05 in both analysis and > 0.9 delta-beta between NeuN+ and Blood and > 0.8 delta-beta between NeuN+ and Lineage tissues (Figure 1D). Four tNGBS assays were designed to the 45 sites.

dnaseCpG-neuron: We contrast NeuN+ cells (n=71; DLPFC and OFC) genome-wide DNA methylation profiles (Illumina, HM450) to PBMC and Lineage genome-wide DNA methylation profiles (Illumina, HM450) using LIMMA separately, identifying 1825 CpG sites that exhibited FDR < 0.05 in both analysis and > 0.8 delta-beta between NeuN+ and Blood (Supplementary Figure 3 & Supplementary Table 12). Differential methylated regions (DMR) were identified by contrasting NeuN+ and Blood genome-wide DNA methylation profiles using bumphunter [1], with significant regions (fwer < 0.05) intersected with significant CpGs to refine 1711 hypermethylated for which GenomicRanges (GR) objects were created +/- 150bp of each CpG (NeuN-GR) for intersection with DNase hypersensitivity GRs (below).

We hypothesized that euchromatic DNA within Blood cells would be preferentially digested by endogenous DNase enzymes and thus provide better targets within for analysis of rare brain derived DNA within cfDNA. To define blood and brain hetero/ euchromatic regions we used DNAse hypersensitivity data produced by the ENCODE consortium (Supplementary Table 3A). We merged the DNase narrowPeaks derived from blood cells (N=23) and brain tissue (N=3) into combined GR objects, Blood-GR and Brain-GR respectively and identified 992 CpG sites (NeuN-GR) that intersected with Blood-GR but not Brain-GR. To reduce the CpG loci, we investigated common euchromatin across the various blood cells by counting number of blood DNase hypersensitivity datasets overlapping each peak ("dataset counts"). Additionally, we calculated the enrichment mean of all blood derived DNase datasets contributing to each Blood-GR narrowPeak ("dataset enrichment"). We identified 67 CpG loci within top $10^{th}$ percentile of the Blood-GR narrowPeaks "dataset counts" and top $10^{th}$ percentile of "dataset enrichment" of which we identified 14 CpG with > 0.5 delta-beta between NeuN+ and Lineage tissues (Supplementary Figure 3B).

Identification of Brain-specific CpH DNA methylation for targeted assay design: Recently, Lister and colleagues illustrated the strikingly unique whole-genome DNA methylation patterns in neurons that, unlike all other human cells, exhibit CpH as their predominant DNA methylation mark [2]. The CpH marks of Neurons (and potentially NeuN-) represent highly unique molecular mark of neurological derived tissue. To refine CpH sites we utilize the WGBS data from FACS sorted (NeuN+ and NeuN-) derived from the Prefrontal Cortex of 1 male and 1 female adult generated by Lister and colleagues (GSE47966).

Smoothing algorithms, such as bumphunter, look for large areas of the genome that display differential methylation between dichotomous features. To maximize our assay specificity/sensitivity for neurological tissue we aimed to identify regions with the highest density of methylated CpH. Using a coverage threshold of 5X we identified 309452 CpH sites that displayed 100% methylation between both male and female samples, whereas only 17765 CpH were found common within NeuN-. We removed X and Y chromosomes and calculated the density of CpH sites +/- 50bp of each hypermethylated CpH site identified. **HighdensityCpH-neuron;** We identified 29336 CpH sites with 1-7 CpH sites within +/- 50bp common to both male and female. Of these, 28 CpH sites had >= 5 CpH sites within a +/- 50bp, of which 8 overlapped, none of which were found within NeuN-, resulting in 20 regions with a high density of CpH DNA methylation specific to NeuN+.

**middensityCpH-neuron;** Additionally, we found 51 CpH sites from the 29336 that had >= 4.5 (average male and female) CpH sites within a +/- 50bp. **CpH-neuronglia;** We also identified 11 CpH sites that were common between NeuN+ and NeuN- that had >= 2 CpH sites within a +/- 50bp. **dnaseCpH-neuron;** Finally we intersected our NeuN+ CpH sites with Blood and Brain chromatin GR (above). Of the 29336 NeuN+ CpH sites we identified 1719 CpH sites within Blood-GR and not Brain-GR. We found 6 CpH sites were found within regions of DNase hypersensitivity sites in over half (>12 of 24) of the blood DNase datasets and which had >2 CpH sites within a +/- 50bp.

*II. Multiplex Bisulfite Sequencing cfDNA Target Regions Assay Design and Testing.*

*Assay Design Rationale:* Genomic DNA sequences ± 5000 bases from the desired target gene regions were acquired from Ensembl and annotated to include exons/introns, the transcriptional and translational start sites, and possible polymorphisms. Assays were designed to interrogate submitted target loci using Pyromark ADSW v1.0 software (Qiagen, Pyrosequencing), in combination with manual assessment. PCR primers were designed to avoid extremely low complexity regions, CpG sites, and high-frequency polymorphisms.

*DNA methylation standards*: CpG DNA methylation standards were purchased from Zymo, CpH DNA methylation standards were constructed from pooled ratios of NeuN+ gDNA and PBMC gDNA. Bisulfite conversion was performed using 100 ng of DNA using the EZ DNA Methylation Kit (Zymo Research) and a modified protocol optimized for low input DNA. 500 µg of resuspended VX Carrier RNA (Qiagen, Cat. # 950280) was added to the M-binding buffer to improve binding efficacy and total yield after treatment. Immediately following the procedure, bisulfite-modified DNA was purified as per the manufacturer's protocol and eluted in 46µL of M-Elution buffer.

*Multiplex PCR Optimization and DNA methylation standards analysis:* In silico designed assays were divided into two groups based on the amplicon size, primer Tm, and GC content, while also avoiding overlapping primer pairs, followed by capillary electrophoresis (CE) of the PCR products using the Agilent 2100 Bioanalyzer system and DNA 1000 chips (Agilent). Based on the QC results, nested PCRs were performed to recover amplicons that did not sufficiently amplify in the initial reaction. In total, three multiplex PCRs and 11 simplex PCRs were generated. PCRs were performed on the DNA methylation controls using the PCR multiplex protocols and thermal cycling conditions reported in supplementary methods. QC of PCR products was performed using either the Agilent 2100 Bioanalyzer or the Qiagen QIAxcel Advanced System.

*Library Preparation and Sequencing*: Prior to library construction, PCR products were pooled and purified using QIAquick PCR Purification Kit columns (Qiagen). Libraries were prepared using the KAPA Library Preparation Kit for Ion Torrent Platforms (Cat# KK8310) and IonXpress™ Barcode Adapters (Thermo Fisher). Next, library molecules were purified using Agencourt AMPure XP beads (Beckman Coulter) and quantified by real-time PCR using the KAPA Library Quantification Kit (Cat.#KK4827). Barcoded samples were then pooled in an equimolar fashion before template preparation was performed on 340 million library molecules using the Ion PGM Templating OT2 200 kit (Thermo Fisher). Following this, enriched, template-positive library molecules were then sequenced on the Ion Torrent PGM sequencer using the Ion PBM™ Sequencing 200 Kit v2 kit with Ion 314™ v2 Chips (Thermo Fisher).

*Sequence Alignment and Data Analysis*: FASTQ files from the Ion Torrent PGM server were aligned to the local reference database using open-source Bismark Bisulfite Read Mapper with the Bowtie2 alignment algorithm. Methylation levels were calculated in Bismark by dividing the number of methylated reads by the total number of reads, considering all CpG sites covered by a minimum of 30 total reads. For CpG methylation assays, an R-square value of >0.9 was required for validation. Each CpH assay had >1 CpH loci with R-square value of >0.75.

III. Measurements of cell-free DNA purity using Weighted Fraction Ratio's (WFR)

Weighted Read Ratio (WRR): Let $r_{s,a}$ be the number of reads for the sample $s = 1, \cdots, S$ and assay $a = 1, \cdots, A$. The weights for assay $a$ was established by dividing ratio of the total reads for assay $a$ $\text{Tot}_a$=sum($r_{s,a}$, for all $s$) to the total reads of all targets Tot=sum($r_{s,a}$, for all $s$ and $a$). The coverage of assay $a$ for sample $s$ can be rescaled to obtain weighted reads (WR)

$$\text{WR}_{s,a} = r_{s,a} / \left( \frac{\text{Tot}_a}{\text{Tot}} \right)$$

This re-scaling is essentially to do row-normalization on matrix formed by the reads $r_{s,a}$.

CfDNA exhibits a well-defined size distribution of around ~180bp, corresponding to nucleosome bound units of DNA (reviewed in [19]). We hypothesized that a ratio of WR between assays >170bp and <170bp could be used as a measure of non-cfDNA contamination within the sample. We investigated the optimal group of assays >170bp to include in this measure by calculating the ratio of summed WRs >170bp (a170) and < 170bp (b170) and performing Students T-test between PBMC (N=10) and cfDNA (N=10) tNGBS results from matched patients (Breachers day 1). This process was repeated, each time removing the shortest assay of the a170. We found the weighted reads difference between PBMC derived gDNA and cfDNA WR of one assay targeting (253bp) and b170 most significant (P= $1 \times 10^{-4}$) (Supplementary Figure 4). We termed this the Weighted Read Ratio (WRR).

$$\text{WRR}_s = \frac{\text{WR}_s(253bp)}{\text{sum}(\text{WR}_{s,a}, a > 170bp)}$$

IV. Laboratory protocols for targeted next generation bisulfite sequencing (tNGBS) of cfDNA

<u>tNGBS protocol steps</u>

A. Cell-free DNA (cfDNA) extraction (1.5 hours)
B. CfDNA quantification (15 mins)
C. Lambda DNA spike-in (15 mins)
D. Bisulfite Conversion (2.5 hours)
E. Sample Dilutions (30 mins)
F. Multiplex Bisulfite PCR's
G. Amplicon pooling
H. Transposase tagging of tNGBS
I. Library Indexing PCR
J. Sample pooling
K. Ampure bead cleanup of library primer-dimer (30 minutes)
L. Quality control KAPA qPCR
M. KAPA qPCR thermal cycling profile
N. Sample Denaturation, PhiX Spike-in and MiSeq loading
O. PCR reaction cocktails and PCR cycling profiles for tNGBS

**A. Cell-free DNA (cfDNA) extraction (1.5 hours)**

i. Perform cfDNA extraction from plasma or serum using the Analytik Jena PME cfDNA extraction kit using the SE/SBS system only.
   a. NOTE: There are two different protocols depending on the starting amount of plasma/ serum (<1ml or 2-5mL)
ii. Elute twice (25uL first and 25uL second) using pre-warmed elution buffer (50uL total).

OPTIONAL STOPPING POINT – Extracted cfDNA may be kept overnight at 4C or frozen at -20C for longer term storage.
NOTE: Minimize freeze-thawing of cfDNA by either quantifying cfDNA (below) prior to freezing OR prior to Lambda spike-in and Bisulfite Conversion.

**B. cfDNA quantification (15 mins)**
Determine the amount of cfDNA extracted by qubit.

i. Use 2uL of each sample with the Qubit High Sensitivity kit.
ii. Calculate the total cfDNA amount remaining after using 2uL for qubit i.e. 48 (remaining vol in uL) X Concentration

**C. Lambda DNA spike-in (15 mins)**
The bisulfite conversion efficiency of each sample is determined by the total conversion of cytosine to thymine of Lambda DNA spiked-into each sample. Lambda DNA is spiked-in at 0.5% W/W into each sample <u>PRIOR</u> to bisulfite conversion (BC). Lambda DNA aliquots at 0.1 or 1ng/uL can be found within the "cfDNA PCR" box within the -20C freezer.

i. Calculate 0.5% of total cfDNA for each sample
ii. Spike in appropriate amount of Lambda DNA, adding ≤ 4 uL to the sample i.e.

**D. Bisulfite Conversion (2.5 hours)**

Bisulfite Conversions (BC) are performed using the MethylCode (Invitrogen) BC kit. NOTE: DO NOT spike-in RNA for samples that have low amounts of cfDNA (RNA spike-in has been found to affect the efficiency of PCRs downstream).

i. Make up the CT conversion reagent by adding the following to the CT conversion reagent powder:
   - 600uL H2O
   - 50uL Resuspension Buffer
   - 300uL Dilution buffer

ii. Vortex the CT conversion reagent for 10 minutes.

iii. Add 100uL of CT conversion reagent to each sample (~50uL)

iv. Elute BC samples in 11uL Elution Buffer.

**E. Sample Dilutions (30 mins)**

To simplify the PCR reactions downstream we dilute samples based on there pre-bisulfite conversion cfDNA amounts.

i. Dilute samples using ddH2O. Do not use Elution Buffer as the EDTA is a chelating agent that dramatically affects bisulfite PCR efficiency.
   - Samples >40ng dilute 4ng/uL
   - Samples 20-40ng dilute to 2ng/uL
   - Samples < 20ng dilute to 1ng/uL

ii. After dilution re-order samples from highest to lowest concentration.

**F. Multiplex Bisulfite PCR's**

The amount of cfDNA extracted from a plasma/ serum sample is often limited. Within each PCR, 4ng or cfDNA is used, therefore it is often not possible to use apply all assays within the tNGBS multiplex. Therefore, the following decision tree (Supplmentary Figure 1) has been created to help researchers decide which PCRs to run.

NOTE: Thermal cycling profiles are labeled the same as the assay names.

i. PCR - Lambda; Two independent PCRs are performed (Lambda 1 and Lambda 3) to amplify Lambda DNA initially pooled within the cfDNA. For each reaction 0.5uL of cfDNA/ Lambda is used. PCR Master mixes can be found at end of document. Also a cheat sheet is available in tNGBS_PCR_cheatsheet.xls. Lambda PCR's must be performed for each sample.

ii. PCR – Primary reactions; PCR Master mix can be found at end of document. Also a cheat sheet is available in tNGBS_PCR_cheatsheet.xls. The first round multiplex PCRs have been optimized for 4ng BC DNA input.

iii. PCR – Nested PCR reactions; PCR Master mix can be found at end of document. It is recommended to use the cheat sheet is available in tNGBS_PCR_cheatsheet.xls. Nested PCR reactions are performed using 1uL of primary PCR from either NGS40_P1/P2 and/ or NGS40_P3

#### G. Amplicon pooling

i. Following PCR amplification of each sample the (up to) 20 tNGBS amplicons/ amplicon pools are mixed together using the ratios in Table 1 below.
ii. Qubit sample and dilute the pooled sample to 3.3ng/uL with DEPC treated $H_2O$.

| Assay/ pool | Pooling (uL) |
|---|---|
| N40_P4 | 0.9 |
| ADS3445 | 0.6 |
| ADS3464 | 0.25 |
| ADS3465 | 0.25 |
| ADS3468 | 1.3 |
| ADS3469 | 0.3 |
| ADS3471 | 0.8 |
| N48_98 | 3.8 |
| ADS3470 | 4.4 |
| ADS3496 | 0.5 |
| ADS3497 | 5.4 |
| ADS3499 | 1.9 |
| N52_09 | 0.8 |
| N52_P1 | 0.25 |
| N52_P2 | 1.1 |
| NS40_P1P2 | 0.8 |
| NS40_P3 | 6 |
| Lambda P1 | 0.25 |
| Lambda P3 | 0.3 |
| WM1&Brain1 | 0.5 |

Supplementary Table 1. PCR pooling ratios.

#### H. Transposase tagging of tNGBS

The cfDNA libraries are generated using the Nextera transposase (Tn5) enzyme supplied within Illumina Nextera® DNA Sample Preparation Kit (FC-121-1031). Ensure the correct kit is used as the Tn5 enzyme from other Nextera products is different.

i. Make Tn5 Master Mix for transposition reaction, multiply by number of samples and add an additional reaction for overhang

| **Tn5 cocktail** | **X1 (5uL)** |
|---|---|
| Tn5 Enzyme | 1 |
| TD Buffer | 2.5 |
| DNA (pooled amplicon) 3.3ng/uL | 1.5 |

Supplementary Table 2. Tn5 Master Mix for transposition reaction.

ii. Incubate at 55C, 5 min and 4C forever

iii. Clean-up reaction immediately using the **MiniElute reaction cleanup kit.** Elute in 10 uL EB buffer.

#### I. Library Indexing PCR

i. Make Indexing PCR Master Mix for each Tn5 transposed sample ensuring each sample has a unique Primer 1 and Primer 2 combination.

| PCR MM | x1 (50uL) |
|---|---|
| Tn5-tNGBS pooled amplicon | 10 |
| H2O | 10 |
| Primer 1 (25uM) | 2.5 |
| Primer 2 (25uM) | 2.5 |
| NebNext PCR MM | 25 |

Supplementary Table 3. Indexing PCR Master Mix.

ii. Thermal cycle as follows:

| | | |
|---|---|---|
| 1 cycle : | 5 min | 72C |
| | 30 sec | 98C |
| 3 cycles: | 10 sec | 98C |
| | 30sec | 63C |
| | 1 min | 72C |

#### J. Sample pooling

i. Samples are pooled (Lo-Bind 1.5mL microcentrifuge tubes) on the basis of the sequencing depth required for each sample i.e. expect 20M reads from MiSeq V2 50PE, therefore 1:20 = 1M reads. It is good practice to use only a portion of each sample for pooling i.e. 10-20uL, just in case the library fails.
ii. Vortex sample and split into <= 200uL (Lo-Bind 1.5mL microcentrifuge tubes) aliquots for Ampure cleanup

#### K. Ampure bead cleanup of library primer-dimer (Time: ~30 minutes)

i. Split pooled library into <200uL and perform Ampure cleanup as described above at 1:1 ratio.
ii. Add equal volume of AMPure XP bead solution to sample within a Low Bind 1.5mL micro centrifuge tube, vortex for 10 seconds, and incubate at RT for 10 minutes. At this ratio, fragments of approximately 100 bp and larger will bind to the beads.
iii. Place the tube on a magnetic stand for 5 minutes, until the solution is clear. With the tube still on the magnetic stand, carefully pipette out and discard the supernatant, leaving behind 5-10 **µ**L so as not to remove any beads. The supernatant contains the unwanted DNA fragments whereas the beads contain the proper size fragments.
iv. Add 500 **µ**L of 80% ethanol to the tube. Immediately remove the tube from the magnetic stand, rotate it 180 degrees and place it back on the magnetic stand. The beads will eventually jump across the ethanol solution, removing any residual binding buffer trapped within the bead pellet. Repeat the rotation 6-10 times. The beads should eventually appear to separate and move across the solution as a cloud instead of a solid bead pellet.

v. With the tube on the magnetic rack, let the beads gather for one minute, then carefully pipette out and discard the supernatant. In this step, the beads should stay against the side of the tube when the ethanol solution is removed completely. Repeat steps 3 and 4 to wash the beads a second time and remove any remaining liquid.

vi. Remove the tube from the magnetic stand and let it sit with the cap open for 5 minutes or until the beads are dry. Small cracks can be observed in the dried bead pellet.

vii. Add **50 µL** of 10 mM Tris-Cl directly to the pellet. Mix at least 15 times using a pipette, let it sit for 10 minutes, then mix again.

viii. Place the tube on the magnetic stand for at least 2 minutes to allow complete capture of the beads. When the suspension is clear, transfer the entire supernatant to a new 1.5 mL tube.

ix. Elute each cleanup into 50uL and combine (usually =100uL)

x. Perform a second round of Ampure cleanup (steps 2-8), eluting library into 20uL 1M Tris-HCl (ph 8.5)

**L. Quality control KAPA qPCR**

A quality control qPCR is performed using the KAPA qPCR Illumina library quantification kit. This qPCR amplifies using illumine sequencing primers and thus measures the amount of "sequencable" library available.

i. Dilute 2uL of library1:100 with DEPC treated water

ii. Dilute 2uL of 1:100 library again to 1:100 = 1:10,000 and

iii. Dilute 10uL of 1:10,000 library 1:10 = 1:100,000

iv. Make KAPA qPCR Master Mix (below) ensuring enough reactions are made for standards. Also we have used PhiX as a positive control previously. Refer to manual for detailed instructions.

| **KAPA qPCR MM** | **X1 (10uL)** |
|---|---|
| KAPA MM + primer | 6 |
| DNA | 2 |
| H2O | 2 |

Supplementary Table 4. KAPA qPCR Master Mix

**M. KAPA qPCR thermal cycling profile**

| | | |
|---|---|---|
| 1 cycle : | 5 min | 95C |
| 35 cycles: | 30 sec | 95C |
| | 45sec | 60C |

**N. Sample Denaturation, PhiX Spike-in and MiSeq loading -** Refer to MiSeq protocol

### O. PCR reaction cocktails and PCR cycling profiles for tNGBS

| NGS40 P1 + P2 | | |
|---|---|---|
| **Component** | **Per 20µl reaction** | **PCR Cycling Conditions:** |
| 10X PCR buffer (Contains 15 mM MgCl2) | 2 µl (1x) | 95C 15 min; 9 x (95C 30 s; 63 - 1C/cycle 30 s; 68C 30 s); 36 x (95C 30 s; 55C 30 s; 68C 30 s); 68C 5 min; 4C □ |
| 25 mM MgCl2 | 1.2 µl (3.0 mM final conc.) | |
| 10 mM dNTPs (2.5 mM each) | 0.4 µl (200 µM of each) | |
| NGS40 P1 Primer Pool | 2 µl | |
| NGS40 P2 Primer Pool | 2 µl | |
| HotStar Taq Polymerase (5 U/µl) | 0.10 µl (0.5 U) | |
| DNA | 3-6 µl of bisulfite treated DNA | |
| Water | Adjust to 20 µl | |

| NGS40 P3 | | |
|---|---|---|
| **Component** | **Per 20µl reaction** | **PCR Cycling Conditions:** |
| 10X PCR buffer (Contains 15 mM MgCl2) | 2 µl (1x) | 95C 15 min; 13 x (95C 30 s; 56 – 0.5C/cycle 30 s; 66C 30 s); 32 x (95C 30 s; 55C 30 s; 66C 30 s); 66C 5 min; 4C □ |
| 25 mM MgCl2 | 1.2 µl (3.0 mM final conc.) | |
| 10 mM dNTPs (2.5 mM each) | 0.4 µl (200 µM of each) | |
| NGS40 P3 Primer Pool | 2 µl | |
| HotStar Taq Polymerase (5 U/µl) | 0.10 µl (0.5 U) | |
| DNA | 3-6 µl of bisulfite treated DNA | |
| Water | Adjust to 20 µl | |

| Lambda BC control PCR | | |
|---|---|---|
| **Component** | **Per 10µl reaction** | **PCR Cycling Conditions:** |
| 10X PCR buffer (Contains 15 mM MgCl2) | 1 µl (1x) | 95C 15 min; 34 x (95C 30 s; 52 30 s; 68C 30 s); 68C 5 min; 4C □ |
| 25 mM MgCl2 | 0.6 µl (3.0 mM final conc.) | |
| 10 mM dNTPs (2.5 mM each) | 0.2 µl (200 µM of each) | |
| Lambda1 & 3 F+R Primer pool (2 uM each) | 1 µl (0.2 uM each final conc.) | |
| HotStar Taq Polymerase (5 U/µl) | 0.05 µl (0.5 U) | |
| DNA | 0.5 µl of bisulfite treated DNA | |
| Water | 6.55 µl | |

| NGS52 P1 | | |
|---|---|---|
| **Component** | **Per 20µl reaction** | **PCR Cycling Conditions:** |
| 10X PCR buffer (Contains 15 mM MgCl2) | 2 µl (1x) | 95C 15 min; 45 x (95C 30 s; 50C 30 s; 68C 30 s); 68C 5 min; 4C □ |
| 25 mM MgCl2 | 1.2 µl (3.0 mM final conc.) | |
| 10 mM dNTPs (2.5 mM each) | 0.4 µl (200 µM of each) | |
| ADS6507 | 0.4 µl | |
| ADS6510 | 0.4 µl | |
| ADS6512 | 0.4 µl | |
| HotStar Taq Polymerase (5 U/µl) | 0.10 µl (0.5 U) | |
| DNA | 2-3 µl of bisulfite treated DNA | |
| Water | Adjust to 20 µl | |

| NGS52 P2 | | |
|---|---|---|
| **Component** | **Per 20µl reaction** | **PCR Cycling Conditions:** |
| 10X PCR buffer (Contains 15 mM MgCl2) | 2 µl (1x) | 95C 15 min; 45 x (95C 30 s; 56C 30 s; 68C 30 s); 68C 5 min; 4C □ |
| 25 mM MgCl2 | 1.2 µl (3.0 mM final conc.) | |
| 10 mM dNTPs (2.5 mM each) | 0.4 µl (200 µM of each) | |
| ADS6506 | 0.4 µl | |
| ADS6508 | 0.4 µl | |
| ADS6511 | 0.4 µl | |
| HotStar Taq Polymerase (5 U/µl) | 0.10 µl (0.5 U) | |
| DNA | 2-3 µl of bisulfite treated DNA | |
| Water | Adjust to 20 µl | |

| NGS52 ADS6509 | | |
|---|---|---|
| **Component** | **Per 20µl reaction** | **PCR Cycling Conditions:** |
| 10X PCR buffer (Contains 15 mM MgCl2) | 2 µl (1x) | 95C 15 min; 45 x (95C 30 s; 53C 30 s; 68C 30 s); 68C 5 min; 4C □ |
| 25 mM MgCl2 | 1.2 µl (3.0 mM final conc.) | |
| 10 mM dNTPs (2.5 mM each) | 0.4 µl (200 µM of each) | |
| ADS6509 | 0.4 µl | |
| HotStar Taq Polymerase (5 U/µl) | 0.10 µl (0.5 U) | |
| DNA | 2-3 µl of bisulfite treated DNA | |
| Water | Adjust to 20 µl | |

| Tn5 indexed adapter PCR | | |
|---|---|---|
| **Component** | **Per 20µl reaction** | **PCR Cycling Conditions:** |
| NEBnext PCR MM | 25 µl | 95C 15 min; 45 x (95C 30 s; 53C 30 s; 68C 30 s); 68C 5 min; 4C □ |
| 25 uM Primer 1 | 2.5 µl | |
| 25 uM Primer 2 | 2.5 µl | |
| DNA (Tn5 pooled amplicon) | 10 µl | |
| Water | 10 µl | |

SUPPLEMENTARY FIGURES

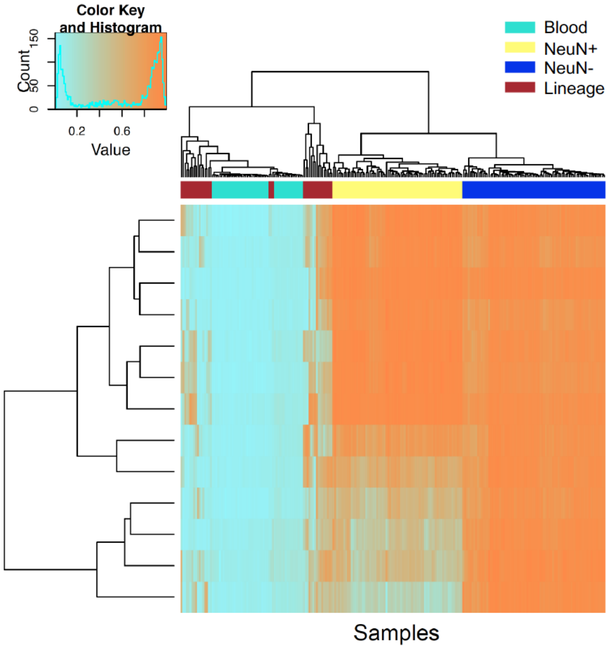


Supplementary Figure 1. Heatmap showing the top 13 probes that were found differentially methylated between NeuN- and blood/ lineage samples.

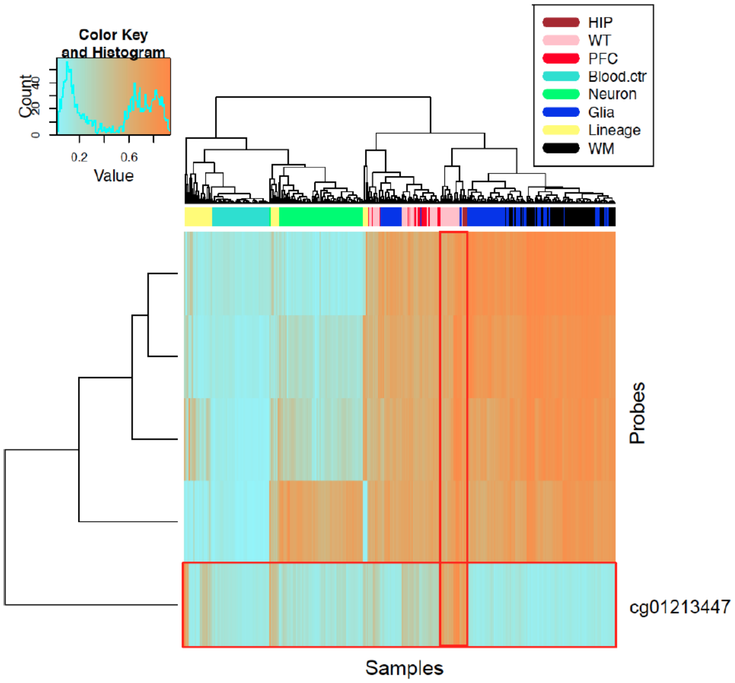


Supplementary Figure 2. Heatmap of DMPs found differentially methylated between hippocampus HIP/ITC and Blood/ Lineage samples and Frontal Cortex whole tissue samples. Probe cg01213447 exhibits hypermethylation within HIP/ ITC samples.

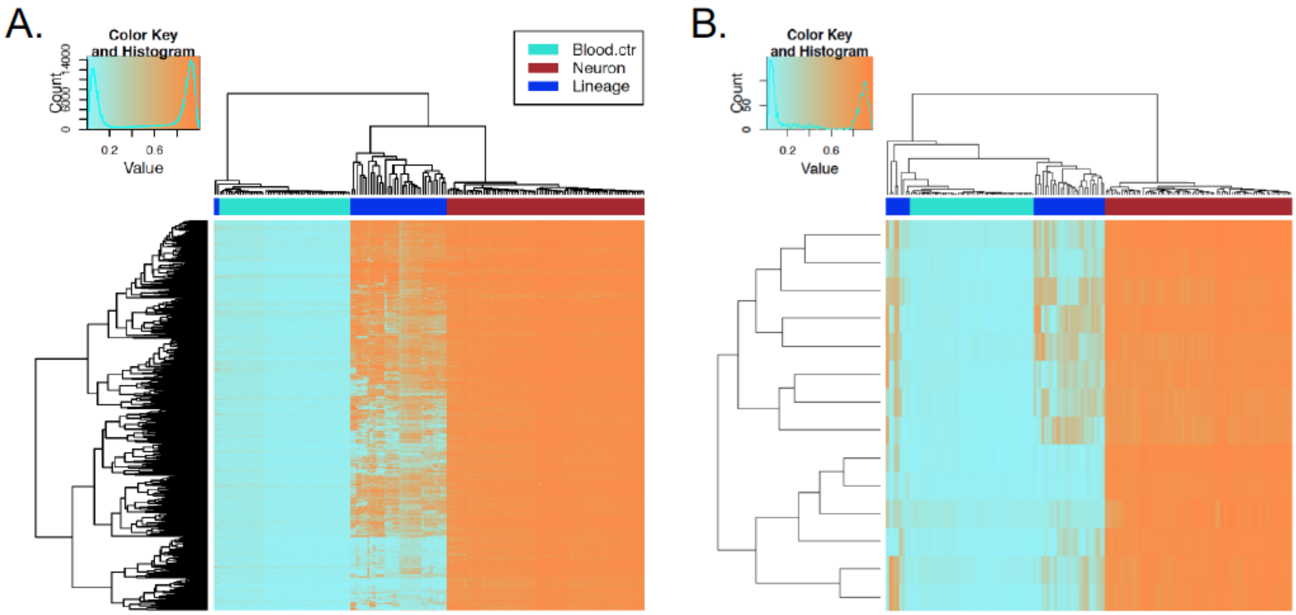


Supplementary Figure 3. A. Heatmap of the NeuN+ hypermethylated CpGs (n=1825) clearly separate the neurons from blood and lineage samples. B. Heatmap of DNA methylation of 14 CpGs found within euchromatic regions of blood cells and heterochromatic regions of brain cells.

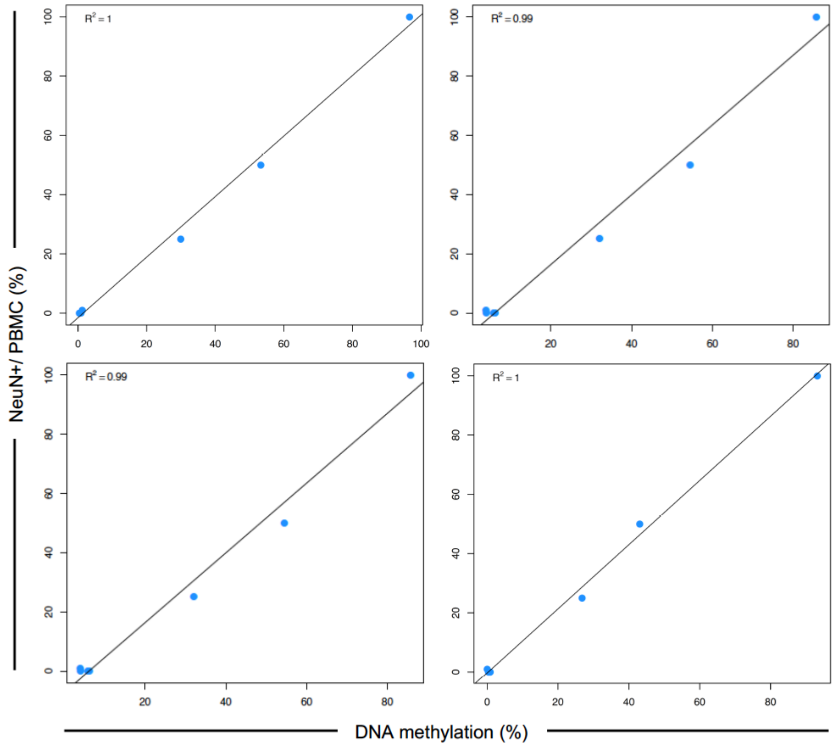


Supplementary Figure 4. Correlation of pooled NeuN+/PBMC (left) and DNA methylation (%) from targeted bisulfite sequencing using assays designed to NeuN+ CpG (left) and CpH (right) methylation.

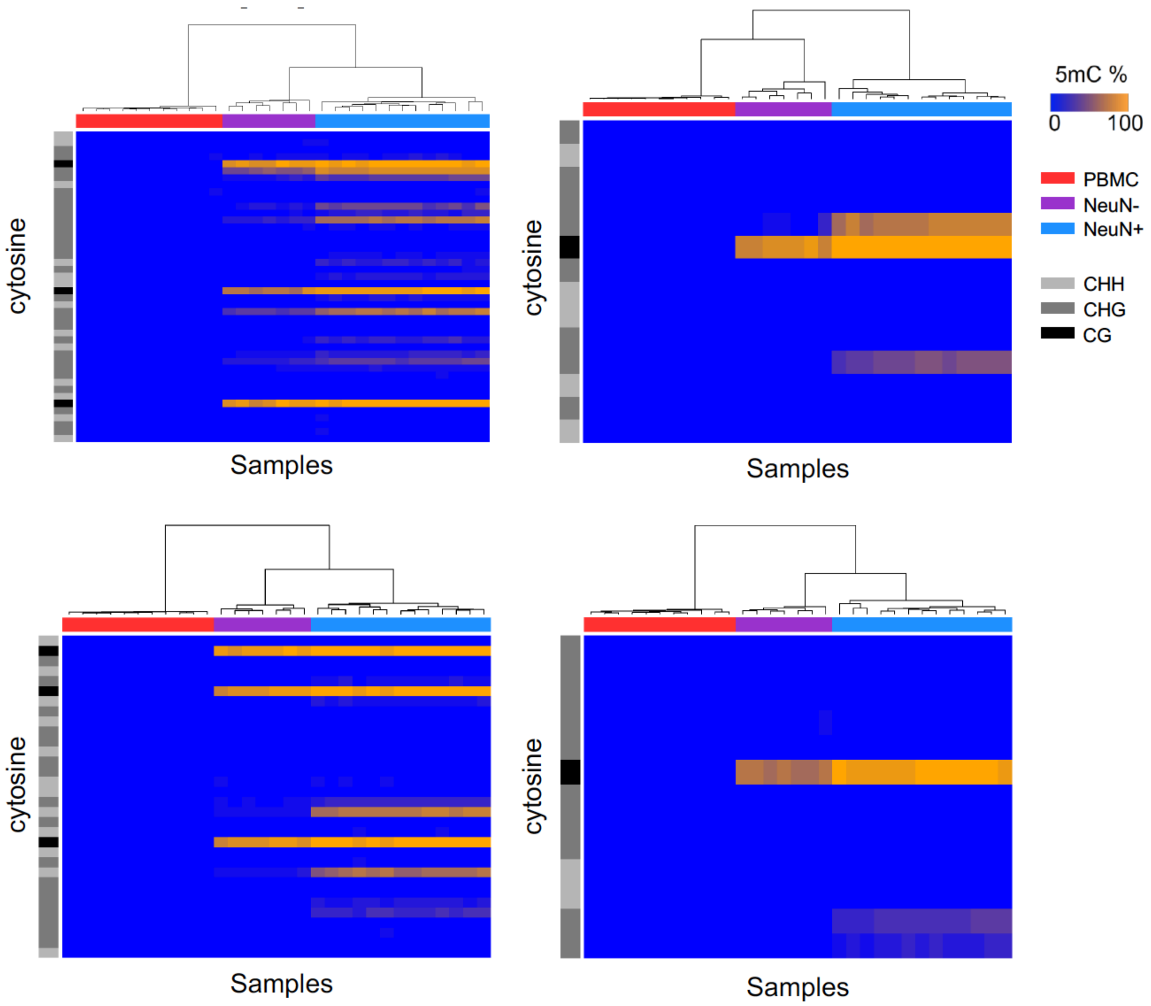


Supplementary Figure 5. Heatmaps showing unsupervised hierarchical clustering of samples analysed by NeuN+ CpG methylation assays. Distinct clustering of DLPFC-NeuN+ cells is driven by CpH methylation whereas clustering of DLPFC-NeuN- cells is driven by CpG methylation.

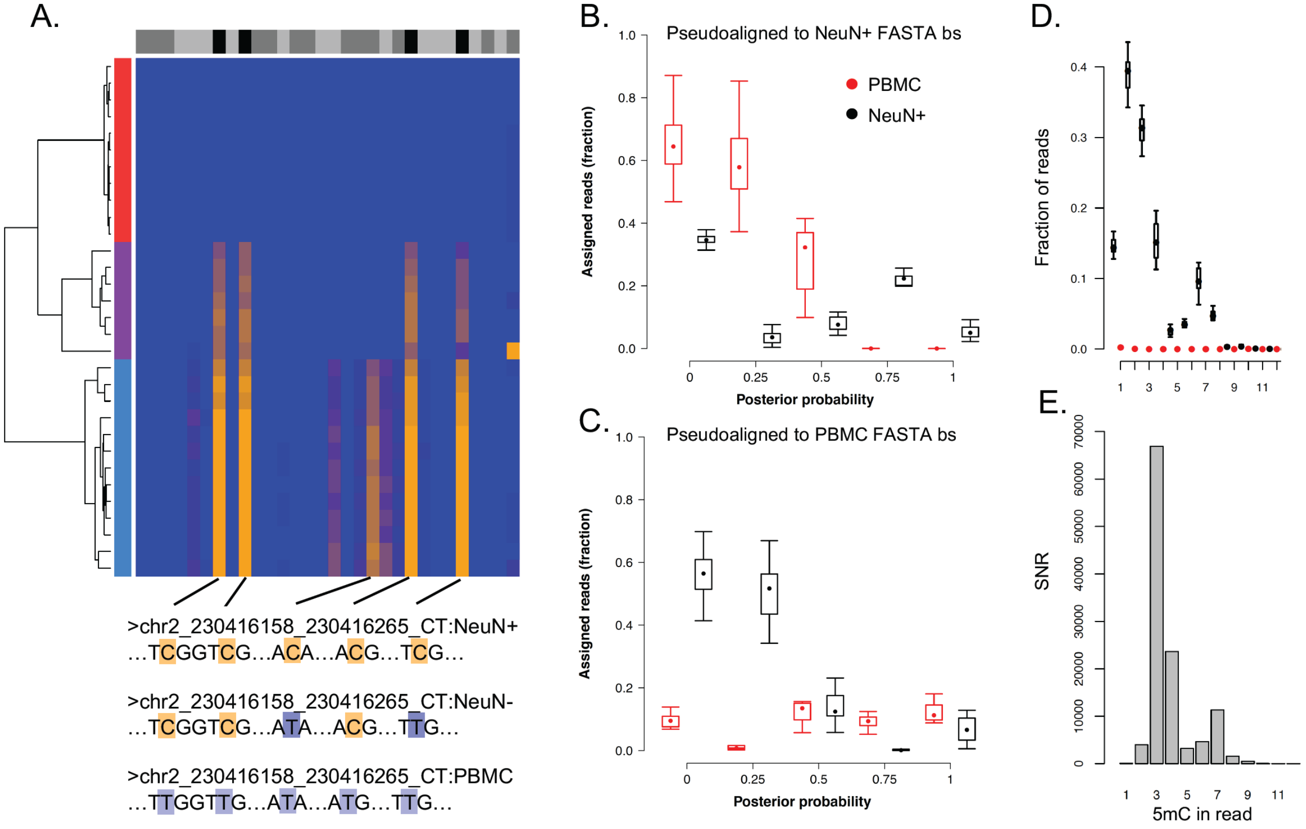


Supplementary Figure 6. K-mer based analysis of bisulfite sequencing reads. A. Example of the binarization of DNA methylation fractions (heatmap) for 3 tissue/cell types (NeuN+, NeuN- and PBMC) into tissue/cell-specific bisulfite FASTA sequences. B. Fraction of pseudoaligned reads to NeuN+ FASTA bisulfite sequence compared to posterior probabilities of pseudoaligned reads. C Fraction of pseudoaligned reads to PBMC FASTA bisulfite sequence compared to posterior probabilities of pseudoaligned reads. D. Example of fraction of bisulfite sequencing reads with 1:N$^{th}$ co-methylation for PBMC and DLPFC-NeuN+. E. A ratio of the fraction of co-methylated counts observed in (D) establishes a signal-to-noise ratio for 1:N$^{th}$ co-methylation events.

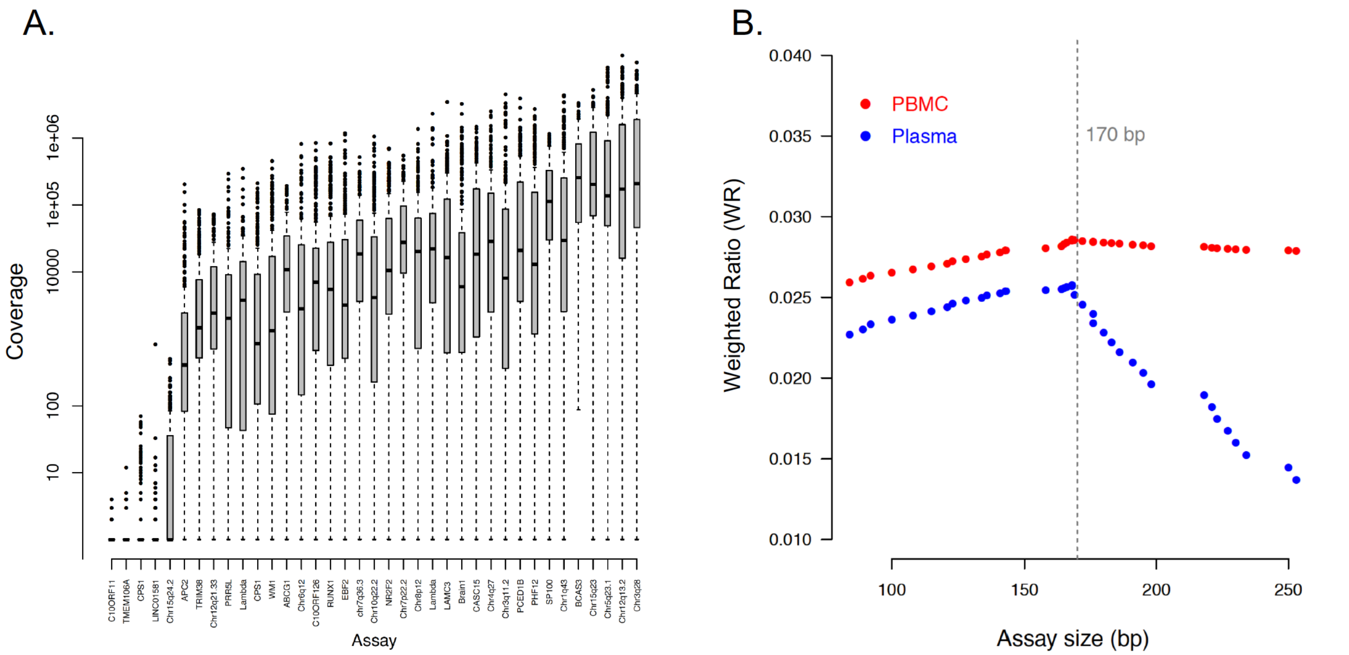


Supplementary Figure 7. A. Coverage of each assay across the population of samples analyzed. B. Weighted Ratio (WR) of each assay by assay length between plasma (cfDNA) and PBMC samples.

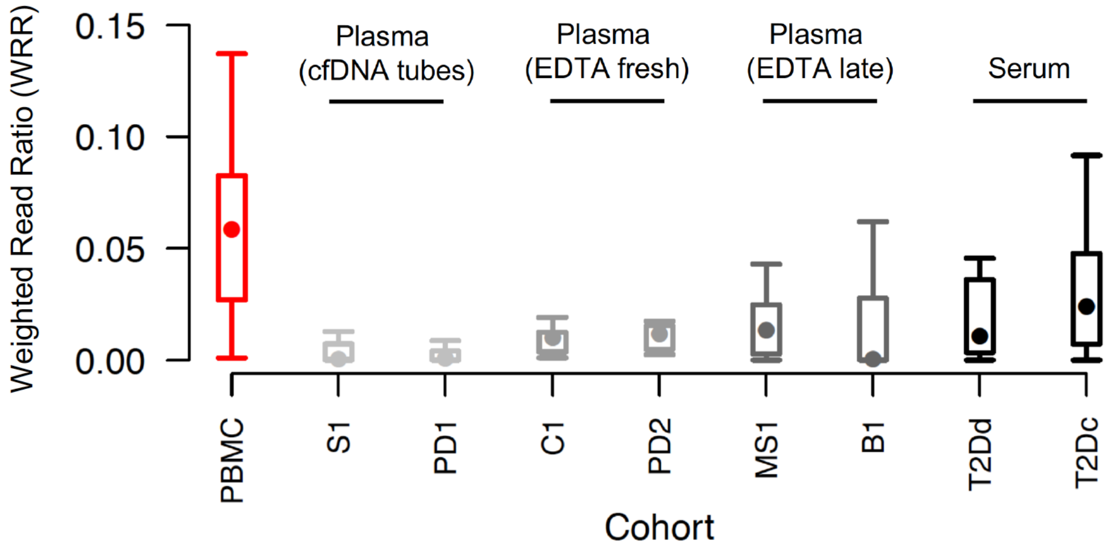


Supplementary Figure 8. Weighted Read Ratio (WRR) across different cohorts with unique sample collection and processing.

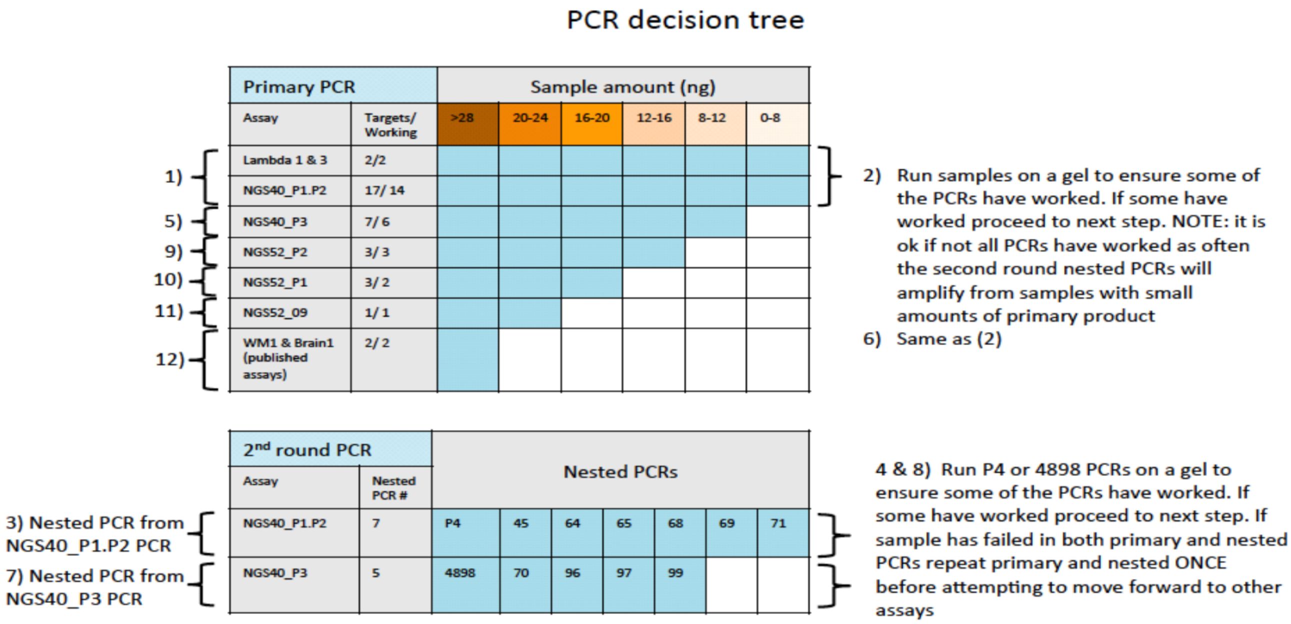


Supplementary Figure 9. PCR decision tree
